# Two PYRIDOXAL PHOSPHATE HOMEOSTASIS PROTEINs are essential for management of the coenzyme in plants

**DOI:** 10.1101/2024.03.03.583161

**Authors:** Peter Farkas, Teresa B. Fitzpatrick

## Abstract

Coenzyme management is believed to be important for the required pool of active enzymes driving metabolic routes to facilitate homeostasis and match environmental circumstance. The coenzyme pyridoxal 5’-phosphate (PLP) (a vitamin B_6_ derivative) is involved in a diverse array of enzyme reactions spanning amino acid to hormone metabolism. However, dedicated proteins that contribute to PLP homeostasis have not yet been studied in plants. Here we demonstrate the importance of proteins annotated PLP HOMEOSTASIS PROTEINs (PLPHPs) for control of PLP in Arabidopsis. A systematic analysis indicates that while most kingdoms have a single *PLPHP* homolog, Angiosperms within the plant kingdom have two. PLPHPs from Arabidopsis bind PLP and exist as monomers in solution in contrast to reported PLP-dependent enzymes from all kingdoms. Disrupting functionality of both homologs perturbs vitamin B_6_ content including a PLP deficit accompanied by impaired and light hypersensitive root growth, unlike biosynthesis mutants. Micrografting studies show that the PLP deficit can be relieved distally between shoots and roots. Yet, supplementation experiments do not restore vitamin B_6_ homeostasis in the absence of PLPHP. A series of chemical treatments probing PLP-dependent reactions, notably those for auxin and ethylene, provide evidence that the physiological role of PLPHPs is dynamic management of PLP. Assays in vitro show that Arabidopsis PLPHP can coordinate both PLP transfer and withdrawal. This study expands our broader knowledge of vitamin B_6_ biology and highlights the importance of PLP coenzyme homeostasis in plants, providing a platform for further investigations in boosting adaptive responses.

**One sentence summary:** PLPHPs contribute to surveillance of vitamin B_6_ homeostasis, likely acting as a rheostat in adaptive responses as a function of the use of the coenzyme PLP.

## INTRODUCTION

Pyridoxal 5ʹ-phosphate (PLP) is an established coenzyme for over 200 reported enzymatic activities, covering 5 of the 6 classes of enzymes, driving amino acid, carbohydrate and lipid metabolism, and is estimated to account for at least 4% of all enzymatic activities (Percudani and Peracchi, 2003). Therefore, PLP is required in every organism for core metabolic processes, the absence of which is fatal. The coenzyme is part of the vitamin B_6_ family that comprises at least six structurally related compounds often referred to as vitamers, specifically pyridoxal (PL), pyridoxine (PN), pyridoxamine (PM) and their phosphorylated derivatives (PLP; PNP, PMP). The pathways for biosynthesis of PLP are well studied including biosynthesis *de novo* pathways in microorganisms and plants, as well as salvage pathways, where other vitamers can be converted into PLP (Liu et al., 2022). The salvage pathways exist in all kingdoms of life including humans, who must take vitamers in their diet. However, PLP, as a coenzyme, is a highly reactive aldehyde capable of forming covalent bonds with amine or thiol groups that can readily damage off-target proteins. For example, excess PLP can lead to formation of unwanted aldimines with exposed epsilon amino (ε-NH_2_) groups of lysines and/or thiazolidine adducts with sulfhydryl groups such as in cysteine, and can thus potentially inactivate proteins or even the chemical role of susceptible metabolites (Hanson et al., 2016). Moreover, regulation of the supply of PLP as a function of demand by enzymes has been proposed to be an important feature of metabolic homeostasis and therefore health and fitness (Liu et al., 2022). Thus, precise control of PLP is required for intact operation of central metabolism. However, management of PLP has received little attention despite its importance. In terms of regulation of levels, early studies suggest negative feedback inhibition of biosynthesis enzymes such as those of *de novo* or salvage pathways that can contribute to homeostasis (Choi et al., 1987; Yang and Schirch, 2000; di Salvo et al., 2011). More recently, an intriguing protein that binds PLP but is without documented enzyme activity has emerged that appears to be involved in controlling the coenzyme (Prunetti et al., 2016; Johnstone et al., 2019; Vu et al., 2020; Ito, 2022). The protein annotated as PLP HOMEOSTASIS PROTEIN (PLPHP) has received considerable attention in humans due to its association with vitamin B_6_ dependent epilepsy, abnormal brain structure, cases of perinatal lethality, and altered neurotransmitter levels when impaired (Darin et al., 2016; Plecko et al., 2017; Johnstone et al., 2019). It has also been studied in microorganisms, and as observed in humans, leads to an amino/keto acid imbalance (Ito et al., 2013; Prunetti et al., 2016; Johnstone et al., 2019). Thus, the protein is associated with pleiotropic phenotypes but its precise physiological functions remain elusive.

In stark contrast to studies in humans (and other animals), metabolism and regulation of PLP has received very limited attention in plants. Studies on the biosynthesis *de novo* and the salvage pathways, albeit relatively recently, have served to illustrate the importance of PLP in plant developmental processes, growth and stress responses (Shi et al., 2002; Tambasco-Studart et al., 2005; Wagner et al., 2006; Colinas et al., 2016; Gorelova et al., 2022; Steensma et al., 2023). Indeed, it has become apparent that balancing of B_6_ vitamers is a crucial feature in plants (Colinas et al., 2016; Steensma et al., 2023), which as sessile organisms assists with the persistent challenge of dealing with environmental variation. For example, an imbalance in B_6_ vitamers can negatively impact nitrogen metabolism, resistance to abiotic and biotic stresses and compromise plant fitness through their roles as antioxidants as well as coenzymes (Liu et al., 2022). However, regulatory mechanisms for PLP homeostasis are ill-defined and PLPHP’s have not yet been studied in plants and thus their role in PLP homeostasis and contribution to growth and development remain unknown.

Here we report on the importance of regulating vitamin B_6_ homeostasis as mediated by *PLPHP* in plants. Interestingly, while most kingdoms have a single *PLPHP* homolog, we show that Angiosperms within the plant kingdom have two, likely resulting from a duplication event before the radiation of monocots and eudicots. Knocking out both homologs in *Arabidopsis thaliana* results in light hypersensitivity of root growth, coincident with impairment of meristem development, suggesting redundancy, and highlighting a role for the proteins in root photomorphogenesis. The absence of both PLPHPs leads to a deficit in PLP levels and perturbation of B_6_ vitamer content throughout the plant, linking their functionality to vitamin B_6_ homeostasis. Grafting studies suggest that a functional deficit in PLP can be relieved distally to restore growth and development in *plphp* mutants. We also provide evidence that the physiological role of PLPHPs is management of PLP and suggest operation through dynamic functioning of PLP-dependent enzymes. Taken together, this study expands our broader knowledge of vitamin B_6_ biology and highlights the importance of PLP homeostasis in plants.

## RESULTS

### Evidence for duplication of PLPHP in Angiosperms

To date only a single homolog of PLPHP has been reported in various kingdoms examined, mainly microorganisms and animals (including humans). As PLPHP has not been studied in plants, we performed a systematics analysis to establish homologs *in plantae*. At the time of the analysis, 8345 genes from 7793 taxa were predicted to be orthologous to the *Escherichia coli* PLPHP homolog (named YggS) (Ito et al., 2013; Prunetti et al., 2016; Tramonti et al., 2022) under the K06997 identifier at the KEGG orthology database (https://www.genome.jp/kegg/ko.html)(Kanehisa et al., 2016). The list included 317 orthologs from 142 plant taxa. Interestingly, more than one ortholog was assigned for over 80% of the green lineage, while over 90% of species in other kingdoms had only one ortholog annotated (Supplementary Table S1). To probe the genetic relation within *plantae*, a phylogenetic tree was constructed based on a multiple sequence alignment performed on 4838 amino acid sequences including 79 sequences from representative species of clades within Archaeplastida (Supplementary Table S2). The phylogenetic tree retraced the evolutionary relation of clades within *plantae* (Figure 1A). Strikingly, core Angiosperms (monocots, eudicots and Ceratophyllales) formed two separate PLPHP clusters, while representatives of basal groups such as Amborellales, Nympheales and Magnoliids had a single sequence with PLPHP orthology (Figure 1A). This suggests that the two homologs of PLPHP (PLPHP1 and PLPHP2) in higher plants result from a duplication event that happened before the radiation of monocots and eudicots (De Bodt et al., 2005).

**Figure 1.**
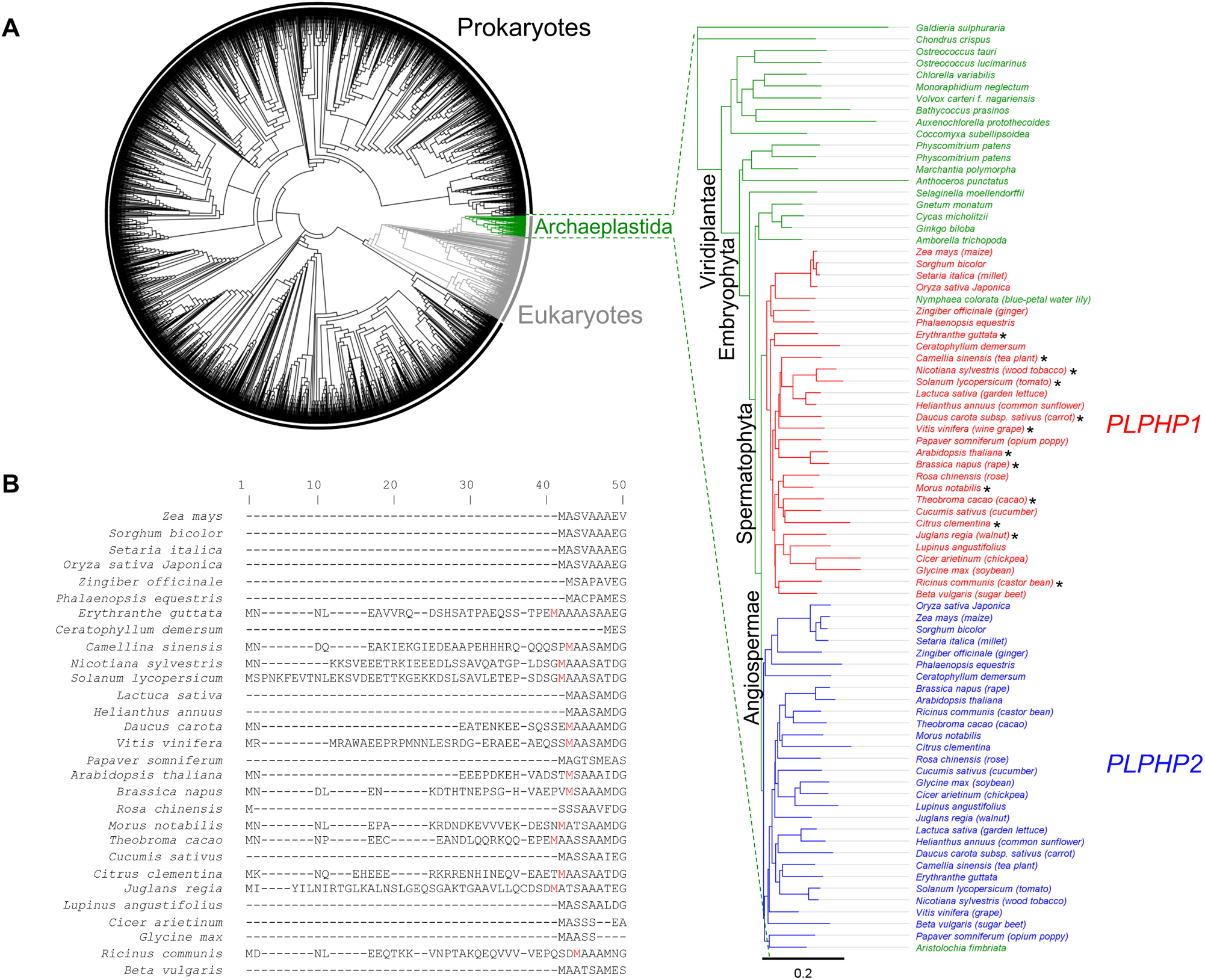
Features of PLPHP amino acid sequences. **A)** Phylogenetic tree of PLPHP representing amino acid sequence similarities of 4838 proteins from eukaryotes and prokaryotes orthologous to YggS from *Escherichia coli*. The branch of Archaeplastida with 79 amino acid sequences is shown in green and the corresponding rooted tree on the right shows PLPHP phylogeny within. Angiosperm sequences (monocots, eudicots and Ceratophyllales) are colored in red and blue corresponding to two putative PLPHP homologs. The scale bar refers to a phylogenetic distance as amino acid substitutions per site. PLPHP1 in certain species is predicted to harbor an N-terminal extension marked with asterisks. **B)** A section of the multiple sequence alignment used in **A** showing the presence of an N-terminal extension in PLPHP1 in certain eudicots.

The Angiosperm PLPHP homologs within a species share 59-79% sequence identity at the protein level. A closer inspection of the sequences within species clearly showed that the sequence of one of the two PLPHP homologs (PLPHP1) in certain eudicots, e.g. Apiaceae (carrot) and Solanaceae (tobacco and tomato) as well as Arabidopsis (Brassicaceae), harbor a short N terminal extension of 15-40 amino acid residues, although apparently clade unspecific (Figure 1A and B). Notably, that of Arabidopsis is confirmed from the recent Arabidopsis PeptideAtlas proteomics resource (van Wijk et al., 2024). By contrast none of the monocot species analyzed harbor this extension.

We conclude that there appear to be two monophyletic groups of PLPHP within Angiosperms of the plant kingdom with a plausible duplication event before the expansion of monocots and eudicots. Furthermore, the presence of an N-terminal extension in PLPHP1 homologs might be a feature inherent to particular members of the eudicots.

### Both PLPHP homologs from Arabidopsis bind PLP in a Schiff-base link as monomers

Similar to PLPHPs characterized from other kingdoms, the plant homologs would be predicted to bind PLP. Indeed, out of the 22 amino acid residues forming the PLP binding pocket (L32, A33, V34, S35, K36, K38, G55, E56, N57, Y58, I82, G83, P84, M164, I166, G203, M204, S205, R220, I221, G222, and T223) in the well-characterized homolog YggS from *E. coli* (Tramonti et al., 2022), 15 showed complete conservation in both homologs of core Angiosperms (Figure 2A). In Arabidopsis, 16 residues are conserved in both homologs, annotated here as AtPLPHP1 (At1g11930) and AtPLPHP2 (At4g26860), which in turn share 71% identity (Figure 2A). To provide evidence that this is actually the case, recombinant protein expression and purification by affinity chromatography was carried out. The purified proteins were pale yellow in color and displayed an absorption peak at 420 nm, distinctive of PLP covalently bound in a Schiff’s base with an active site lysine, under the conditions used (Metzler et al., 1982) (Figure 2B). AtPLPHP1 could be purified in much higher amounts compared to AtPLPHP2 and was used for further tests. Treatment with sodium borohydride caused a hypsochromic shift of the absorption peak in AtPLPHP1 to 320 nm consistent with reduction of the Schiff’s base (Zito and Martinez-Carrion, 1980) (Figure 2B). Furthermore, treatment with sodium hydroxide caused a hypsochromic shift of the absorption peak at 420 nm to 388 nm in line with the release of PLP (Ghatge et al., 2012) (Figure 2B). The bound chromophore was subjected to HPLC and had the same retention time as a commercial standard of PLP (Figure 2C). The predicted active site lysine involved in Schiff’s base formation is K55 in AtPLPHP1 (Figure 2A). Expression and purification of the AtPLPHP1 K55A mutant followed by spectral analysis indicated an at least 10-fold lower amount of bound PLP, thus supporting the involvement of K55 in Schiff’s base formation with the coenzyme (Figure 2B). Although, it should be noted that a second lysine not involved in Schiff’s base formation with PLP but required for functionality as demonstrated with the *E. coli* homolog (Tramonti et al., 2022) is conserved in the Arabidopsis orthologs (Figure 2A).

**Figure 2.**
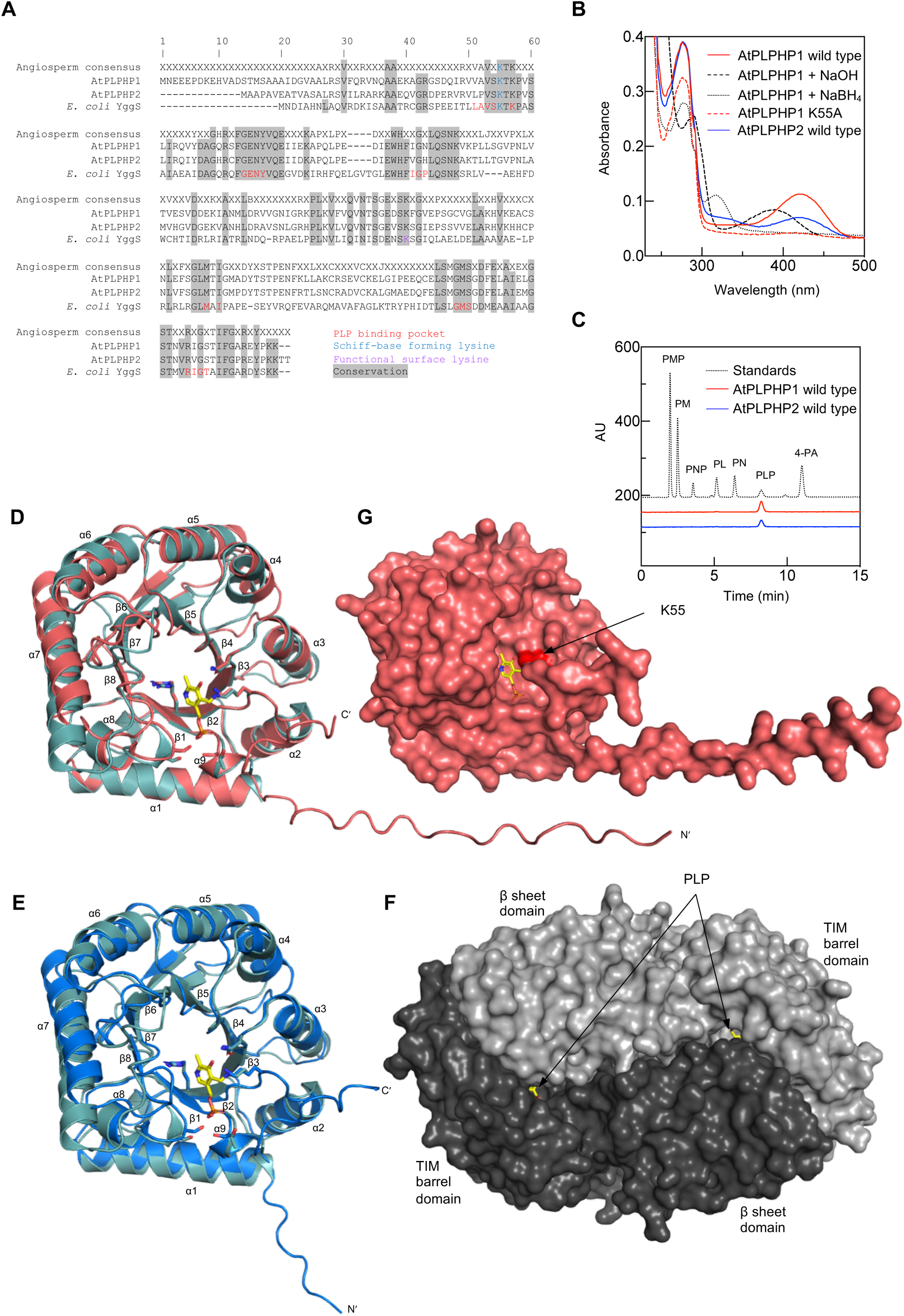
Features of PLPHP1 and PLPHP2 from Arabidopsis. **A)** Angiosperm consensus sequence showing residues that are conserved in at least 95% of the Angiosperm species analyzed. AtPLPHP1 and AtPLPHP2 refer to the two paralogs in *Arabidopsis thaliana*, while YggS is the *Escherichia coli* ortholog. Conserved residues are highlighted in gray. Residues forming the PLP binding pocket in the *E. coli* ortholog are indicated in red, the lysine residue forming a covalent bond with PLP is indicated in blue, the functionally important non-PLP binding lysine residue in the *E. coli* ortholog is indicated in purple. **B)** Absorption spectrum of AtPLPHP1 (red), AtPLPHP2 (blue), AtPLPHP1_K55A (red dashed line) in 50 mM HEPES buffer at pH 7.4 in the presence of 300 mM NaCl. In each case, 75 μM of protein was analyzed. The black dashed and dotted lines represent the same AtPLPHP1 sample treated with sodium hydroxide (NaOH) or sodium borohydride (NaBH_4_). **C)** HPLC chromatograms of purified AtPLPHP1 (red) or AtPLPHP2 (blue) and standards as indicated (black dotted line). **D-E)** Alphafold models of AtPLPHP1 (red cartoon) and AtPLPHP2 (blue cartoon) superposed with the X-ray crystal structure of *E. coli* YggS (turquoise cartoon) (PDB:1w8g). **F)** Surface topology of the *Geobacillus stearothermophilus* alanine racemase homodimer with individual monomers shaded in gray (PDB:1sft). PLP molecules are shown in yellow. **G)** Surface topology of the Alphafold model of AtPLPHP1 from **D** with the surface exposed Schiff-base forming lysine residue highlighted in red and a PLP molecule inserted into the putative PLP binding pocket.

A theoretical structural 3D model of either AtPLPHP1 or AtPLPHP2 generated with Alphafold (Jumper et al., 2021) predicts an eight stranded parallel β-barrel surrounded by eight α-helices (β_8_α_8_), i.e. the triose phosphate isomerase (TIM)-barrel fold (Wierenga, 2001) (Figure 2D, E). Structural superposition of the predicted 3D models with the solved X-ray structure of *E. coli* YggS (Tramonti et al., 2022) indicates conservation of the overall topology (Figure 2D, E). The TIM-barrel fold is one of the most abundant and versatile topologies and a typical feature of Type III PLP-dependent proteins, such as ornithine decarboxylase or alanine racemase (Eliot and Kirsch, 2004). However, Type III PLP-dependent enzymes are two-domain proteins comprising the TIM-barrel at the N-terminus and a C terminal β-sheet module (Eliot and Kirsch, 2004) (Figure 2F). Members of this class form homodimers in a head-to-tail configuration such that the β-sheet module shields the bound PLP molecules within the TIM-barrel of the opposing monomer and contributes to the active site found at the interface of both monomers (Figure 2F). Both AtPLPHP1 and AtPLPHP2 lack the characteristic C terminal β-sheet module present in the Type III fold enzymes that is responsible for these features. Furthermore, the lysine that is involved in binding PLP appears to be solvent exposed (Figure 2G), similar to its *E. coli* ortholog (Ito et al., 2013). Interestingly, gel-filtration analysis demonstrated that both AtPLPHP1 and AtPLPHP2 behave as monomers (estimated molecular mass and theoretical monomer mass 28.8/27.8 kDa (AtPLPHP1) and 27.7/26.6 kDa (AtPLPHP2), respectively) (Supplementary Figure S1).

Thus, both of the Arabidopsis PLPHPs have the capacity to bind PLP. They behave as monomers with PLP likely solvent exposed similar to the reported oligomeric state of bacterial and yeast homologs (Eswaramoorthy et al., 2003; Prunetti et al., 2016; Tremiño et al., 2017; Tramonti et al., 2022), although the human ortholog is reported to exist as both monomers and dimers *in vitro* (Tremiño et al., 2018; Fux and Sieber, 2020). Significantly, the monomeric state of Arabidopsis PLPHPs contrasts with reported PLP-dependent enzymes from all kingdoms, all of which have higher oligomeric states (Eliot and Kirsch, 2004).

### The absence of PLPHPs leads to light hypersensitive root tip developmental defects

In order to investigate the functional requirement of the PLPHPs in Arabidopsis, we obtained lines SAIL_675_A03 and GK-740_A10 corresponding to T-DNA insertions in the coding sequences of AtPLPHP1 and 2, respectively (Figure 3A). The lines were isolated to homozygosity for the T-DNA insertion and reverse transcriptase quantitative PCR (RT-qPCR) indicated reduced expression of the genes and were therefore annotated as *plphp1-1* and *plphp2-1*, respectively (Figure 3A and B). We also availed of the Clustered Regularly Interspaced Short Palindromic Repeats (CRISPR)-Cas9 technology to generate knockout mutants of the individual PLPHPs in Arabidopsis. Specifically, we assembled constructs containing three single guide RNA sequences targeting either *AtPLPHP1* or *AtPLPHP2* driven by the *U6* promoter, in the RU051 vector that contains *SpCas9*, expression of which is driven by the parsley *PcUBI4-2* promoter, as well as a red fluorescent protein driven by the seed specific promoter of *OLE1* (At4g25140) as a plant selection marker (Ursache et al., 2021). The T1 generation of plants were screened for red fluorescence combined with PCR analysis and allowed us to select plants carrying deletions in *AtPLPHP1* or *AtPLPHP2*. In the T2 generation, we could isolate transgene-free plants and confirmed the presence and the position of the deletions by sequencing (Figure 3A; Supplementary Figure S2). Reduced expression of the genes was confirmed by RT-qPCR and the lines were annotated as *plphp1-2* and *plphp2-2*, respectively (Figure 3A and B). To confirm altered expression at the protein level, we attempted to raise polyclonal antibodies against the individual AtPLPHP1/2 recombinant proteins. The antibody against AtPLPHP1 recognized this protein and showed no detectable expression of *AtPLPHP1* in *plphp1-1* and *plphp1-2*, whereas it was detected in *plphp2-1* and *plphp2-2* (Figure 3C). However, despite several attempts, we were unable to successfully immunochemically detect AtPLPHP2.

**Figure 3.**
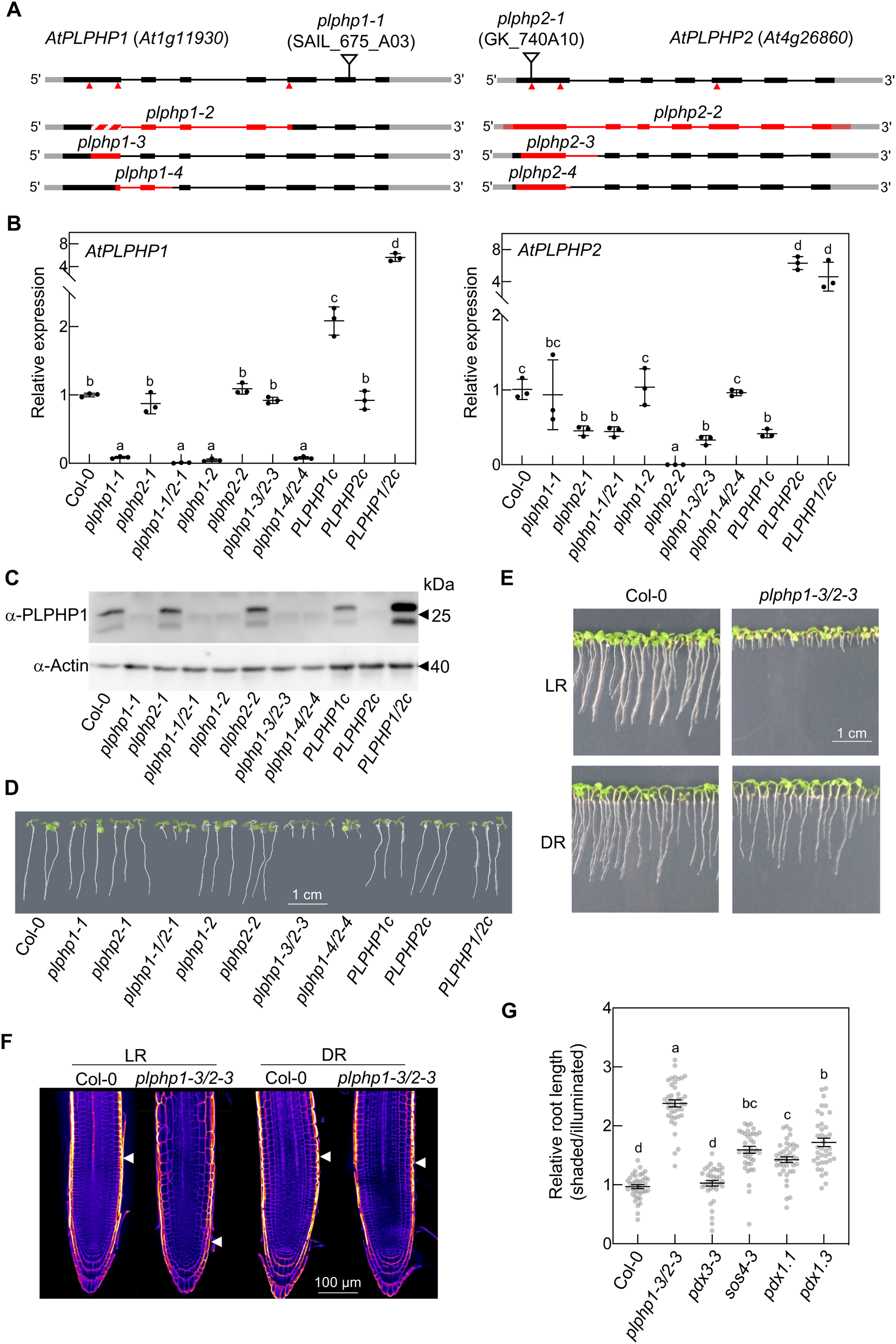
Study of *PLPHP* mutants in Arabidopsis. **A)** Schematic representation of locus At1g11930 (left) and At4g26860 (right) with the position of T-DNA insertions (top) and CRISPR generated deletions (bottom). Black and gray rectangles represent exons and untranslated regions, respectively. Red arrowheads mark CRISPR target sites. The hashed red coloring represents a genetic inversion in *plphp1-2*. **B)** Transcript abundance of *AtPLPHP1/2* in various lines relative to the endogenous control gene *PP2AA3* normalized to expression in Col-0 (wild type). The letters represent significantly different groups (p≤0.05) according to a Brown-Forsythe ANOVA test with Welch’s correction. **C)** Immunochemical detection of AtPLPHP1 protein in crude extract (20 μg) from 16-days-old Arabidopsis lines grown on soil under a 16-hour photoperiod (120 μmol photons m^−2^ s^−1^) at 22°C and 8 hours dark at 18°C. The same blot probed with a monoclonal α-actin antibody is shown below. Black arrows mark the position of molecular mass markers. **D)** Root phenotype of *AtPLPHP1/2* mutant lines and complementing lines 5 days after germination grown on vertical culture plates under a 16-hour photoperiod (120 μmol photons m^−2^ s^−1^) at 22°C and 8 hours dark at 18°C. **E)** Development of Col-0 and *plphp1-3/2-3* seedlings on vertical culture plates with the root area illuminated (top; LR) or shaded (bottom; DR) at 6 days after germination under a 16-hour photoperiod (120 μmol photons m^−2^ s^−1^) at 22°C and 8 hours dark at 18°C. **F)** Representative microscope images of Col-0 and *plphp1-3/2-3* root tips at 6 days after germination under a 16-hour photoperiod (120 μmol photons m^−2^ s^−1^) at 22°C and 8 hours dark at 18°C. Plants were grown on vertical culture plates with the root area illuminated (left; LR) or shaded (right; DR). Seedlings were cleared with ClearSee and cell walls were stained with Calcofluor white. White arrows indicate the first cortical cell with a length at least equal to its width marking the end of the meristematic zone. **G)** Length of shaded roots of indicated plant lines relative to the average length of illuminated roots of the same lines at 6 days after germination. Data represent the mean of 36-40 biological replicates ± standard error. Letters represent significantly different groups (p≤0.05) according to an ordinary one-way ANOVA with Tukey’s multiple comparison test.

With these lines in hand (see Supplementary Table S3 for the list of lines), we next checked for a morphological phenotype but no obvious defect was observed throughout the plant life cycle when grown on soil under standard laboratory conditions (Supplementary Figure S3). We surmised that the Arabidopsis PLPHP proteins may be redundant and thus crossed the T-DNA insertion lines *plphp1-1* and *plphp2-1* to generate the *plphp1-1 plphp2-1* double mutant (Figure 3B and C). We also targeted both Arabidopsis *PLPHP* genes simultaneously using CRISPR/Cas9 to generate double mutants employing three target sites per gene as above and isolated two independent lines with lesions in both homologs that were annotated as *plphp1-3/4 plphp2-3/4* (Figure 3A-C; Supplementary Figure S2; Supplementary Table S3). No morphological phenotype allowed us to distinguish any of these lines from wild type when grown on soil under standard laboratory conditions (Supplementary Figure S3). However, growth of the double mutant lines on sterile culture plates demonstrated a clear stunting of root growth in the double mutant lines *plphp1-1 plphp2-1* and *plphp1-3/4 plphp2-3/4* (Figure 3D). As the double mutant lines behaved similarly, a more detailed analysis was carried out on *plphp1-3 plphp2-3*. Initially, we assessed key stages of germination and seedling development which showed that *plphp1-3/2-3* germinated as for wild type but was retarded in root development thereafter (Supplementary Figure S4A and B). Notably, there was no significant difference in cotyledon or leaf area of the mutant seedlings compared to wild type (Supplementary Figure S4C and D). Thus, the defect in *plphp1/2* appeared to be localized to the root. Importantly, reintroduction of either *AtPLPHP1* or *AtPLPHP2* or both into *plphp1-3/2-3* with expression driven by the respective upstream regions and similar or above that of wild type (Figure 3B and C), completely rescued the root developmental phenotype (Figure 3D; Supplementary Figure S5A). These lines were therefore regarded as complementing lines and annotated *PLPHP1c*, *PLPHP2c*, and *PLPHP1*/*2c*, respectively (Figure 3B-D; Supplementary Table S3). As seedlings were entirely exposed to light in the above sterile culture plate experiments and light has been shown to impact root development (Cabrera et al., 2022), we next shaded the root area under which root growth was considerably improved in the *plphp1-3/2-3* double mutant, although not completely (Figure 3E; Supplementary Figure S5B). A more detailed microscopic analysis of the root tip in *plphp1-3/2-3* upon light exposure indicated a severely perturbed architecture (Figure 3F). In particular, the meristem size is severely reduced (defined by the number of cortex cells in a file extending from the quiescent center to the first elongated cell (for which the length is double the width in the median, longitudinal section)) (Figure 3F). Shading of the roots restores the meristem size of *plphp1-3/2-3* to that of wild type (Figure 3F). To assess the specificity of this phenotype, we examined other vitamin B_6_ biosynthesis mutants, *pdx1.1*, *pdx1.3*, and *sos4.3* that have been reported to have short root phenotypes (Boycheva et al., 2015; Gorelova et al., 2022), although the effect of shading was not tested in these studies. Interestingly, while growth of the roots of these mutants was slightly improved upon shading compared to illumination, it was less pronounced than with *plphp1-3/2-3* (Figure 3G), suggesting that mutation of *PLPHP* renders Arabidopsis seedlings particularly sensitive to root illumination.

Taken together, the data suggests that AtPLPHP1 and AtPLPHP2 are functionally redundant. Their absence leads to light hypersensitive root developmental defects that can be alleviated by shading or completely by the reintroduction of At*PLPHP1* or *2*. Thus, it appears that the presence of AtPLPHPs are crucial to minimize growth and developmental defects during root photomorphogenesis.

### AtPLPHPs contribute to B_6_ vitamer homeostasis and maintenance of PLP levels

Next, to assess the contribution of AtPLPHP1/2 to vitamin B_6_ homeostasis in Arabidopsis, we profiled all vitamers using an established protocol (Steensma et al., 2023). Profiling of fully illuminated whole seedling material indicated considerably lower amounts of PLP in the *plphp1-3/2-3* double mutant line compared to wild type (Figure 4A). On the other hand, PMP was not significantly changed (Figure 4A). Interestingly, whereas PNP could not be detected in wild type, it accumulated in *plphp1-3/2-3* (Figure 4A). Furthermore, of the non-phosphorylated vitamers, PL was decreased in *plphp1-3/2-3* mutant lines, whereas PN and 4-pyridoxic acid (4-PA) were increased (Figure 4A). Notably, 4-PA is a degradation product of PLP and PL (Yagi et al., 1983; Fukuwatari et al., 2008). Significantly, the alteration of B_6_ vitamers in *plphp1-3/2-3* was largely reversed in the complementing lines (i.e. *plphp1-3/2-3* carrying either AtPLPHP1, AtPLPHP2 or both), with the exception that the PLP content remained low in the PLPHP2c line (Figure 4A). The restoration of the B_6_ vitamer profile was not observed in transgenic lines which carried a copy of either AtPLPHP1 or AtPLPHP2 in which the active site lysine residue that forms the Schiff-base linkage with PLP was mutated (Figure 4A). We can therefore conclude that PLPHPs are required for maintenance of vitamin B_6_ balance in Arabidopsis, which furthermore relies on the active site lysine residue.

**Figure 4.**
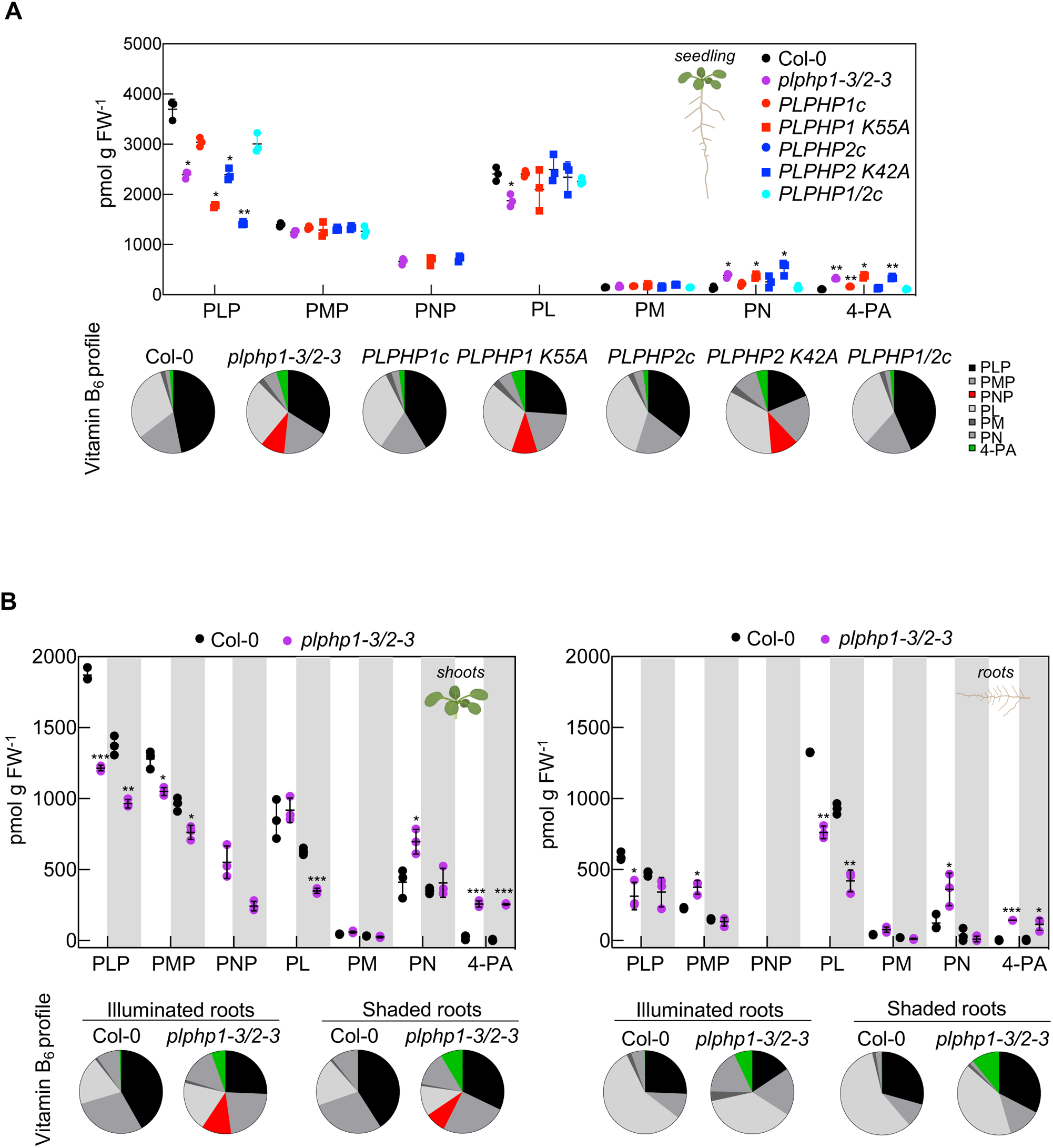
Vitamin B_6_ profiling of lines. **A)** Scatter dot plot (top) showing the abundance of seven vitamin B_6_ related compounds (pyridoxal 5ʹ-phosphate (PLP), pyridoxamine 5ʹ-phosphate (PMP), pyridoxine 5ʹ-phosphate (PNP), pyridoxal (PL), pyridoxamine (PM), pyridoxine (PN), 4-pyridoxic acid (4-PA)), in 6 days old seedlings of various lines as indicated (Col-0 is the wild type, *plphp* double mutant *plphp1-3/2-3*, and the latter carrying either the *PLPHP1* wild type, *PLPHP1 K55A* mutant, *PLPHP2* wild type, *PLPHP2 K42A* mutant or *PLPHP1* and *PLPHP2* wild type transgenes, respectively) grown with full illumination on vertical culture plates under a 16-hour photoperiod (120 μmol photons m^−2^ s^−1^) at 22°C and 8 hours dark at 18°C. PNP was only detected in the lines as indicated, so statistical analysis was omitted for this compound. Letters represent significantly different groups (p≤0.05) according to an ordinary one-way ANOVA with Tukey’s multiple comparison test. The same results are represented as pie charts on the bottom showing the relative distribution of the individual vitamers. **B)** Scatter dot plot (top) showing the abundance of vitamin B_6_ related compounds as in **A** in separated shoots and roots of 6 days old Col-0 (wild type) and *plphp1/2* seedlings grown as in **A** but with roots either illuminated (white) or shaded (gray). PNP could not be detected in Col-0 so statistical analysis was omitted for this compound. Statistical analysis was performed by unpaired *t*-tests with Welch’s correction between genotypes within the same conditions (*p≤0.05,**p≤0.005, ***p≤0.0005, ****p≤0.0001). The same results are represented as pie charts on the bottom showing the relative distribution of the individual vitamers.

To associate the root developmental defect observed upon illumination with vitamin B_6_ content, we next profiled the vitamers in separated shoot and root material when the roots were either exposed to illumination or shaded. Focusing on shoot tissue first, the most pronounced change was decreased PLP (and PMP albeit less pronounced) in the *plphp1-3/2-3* double mutant compared to wild type and regardless of the roots being illuminated or shaded (Figure 4B). Interestingly, the shoot PLP and PMP content was lower when roots were shaded (Figure 4B). PNP was readily detected in the mutant shoots but not in those of wild type (Figure 4B). For the non-phosphorylated vitamers, PL contents were decreased in the shoots of *plphp1-3/2-3*, but only when roots were shaded, PM levels were unchanged in either condition and PN levels were slightly increased when roots were illuminated (Figure 4B). The 4-PA levels were substantially increased in *plphp1-3/2-3* shoots compared to wild type regardless of the roots being illuminated or shaded (Figure 4B). With regards to the roots, the PLP content was decreased significantly in *plphp1-3/2-3* compared to wild type when roots were illuminated but not under shading (Figure 4B). Moreover, root PLP content was much lower than that of the shoot (Figure 4B). PMP levels were slightly higher in illuminated roots of *plphp2*, PNP could not be detected in roots. There was a substantial decrease in PL in *plphp1-3/2-3* under either shading or illumination, PMP did not change and PN was increased in the illuminated tissue (Figure 4B). As for shoots, 4-PA levels were increased in the roots of the *plphp1-3/2-3* double mutant under either shading or illumination (Figure 4B).

We conclude that disruption of PLPHPs in Arabidopsis triggers an overall deficit in PLP and a general perturbance of other B_6_ vitamer levels, characterized at the tissue level by detection of PNP in the shoots and a deficit of PL in the roots. Therefore, PLPHP’s are essential for maintenance of PLP levels and for B_6_ vitamer homeostasis in Arabidopsis. Moreover, the increase in 4-PA in both tissues of the *plphp1-3/2-3* mutant suggests that PLPHPs protect PLP from degradation. Furthermore, it can be concluded that shoot tissue development is recalcitrant to the perturbance in B_6_ vitamer status in *plphp1-3/2-3* but the changes render the root tissue hypersensitive to illumination resulting in severe morphological defects.

### Evidence for distal servicing of PLP levels

Although PLP was deficient in shoots as well as in roots of *plphp1-3/2-3* lines, yet no morphological phenotype was observed in the shoot, we ascertained that a deficit in PLP/PL in *plphp1-3/2-3* might be the most likely explanation for the root hypersensitivity to light. Therefore, we next attempted to rescue the phenotype of *plphp1-3/2-3* by supplementation with non-phosphorylated B_6_ vitamers (PL, PN, PM). These vitamers can be added to the culture plate media and are taken up by the roots, which then due to the ubiquitous presence of salvage pathway enzymes such as the PL/PN/PM kinase SOS4 and pyridox(am)ine phosphate oxidase PDX3 in Arabidopsis (Shi et al., 2002; Gorelova et al., 2022; Steensma et al., 2023) could be expected to service both the PLP and PL deficit. Although the stunted root phenotype of *plphp1-3/2-3* was alleviated under root illumination in the presence of PL, PN or PM (Figure 5A; Supplementary Figure 5C), complete rescue was not observed, and is in contrast to the full rescue observed with the genetically complemented lines (Figure 3D). In line with this, the B_6_ vitamer diagnostic features of *plphp1-3/2-3* persisted upon supplementation (decreased PLP in shoots and roots, increased 4-PA, increased PNP in shoots, decreased PL in roots) (Figure 5B). This is in contrast to what has been reported for biosynthesis *de novo* mutants, e.g. *pdx1*, which are deficient in PLP and where supplementation with the non-phosphorylated vitamers rescues developmental defects (Titiz et al., 2006; Wagner et al., 2006). Notably, mutants of the PLP biosynthesis *de novo* (and salvage pathways) have general developmental impairments in both shoots and roots as observed in soil and culture plates (Shi et al., 2002; Titiz et al., 2006; Wagner et al., 2006; Gorelova et al., 2022; Steensma et al., 2023). Thus, *plphp1-3/2-3* is distinct from previously characterized vitamin B_6_ mutants in phenotype and in terms of servicing by vitamer rescue suggesting that PLPHP itself is vital for vitamin B_6_ homeostasis and conditional development.

**Figure 5.**
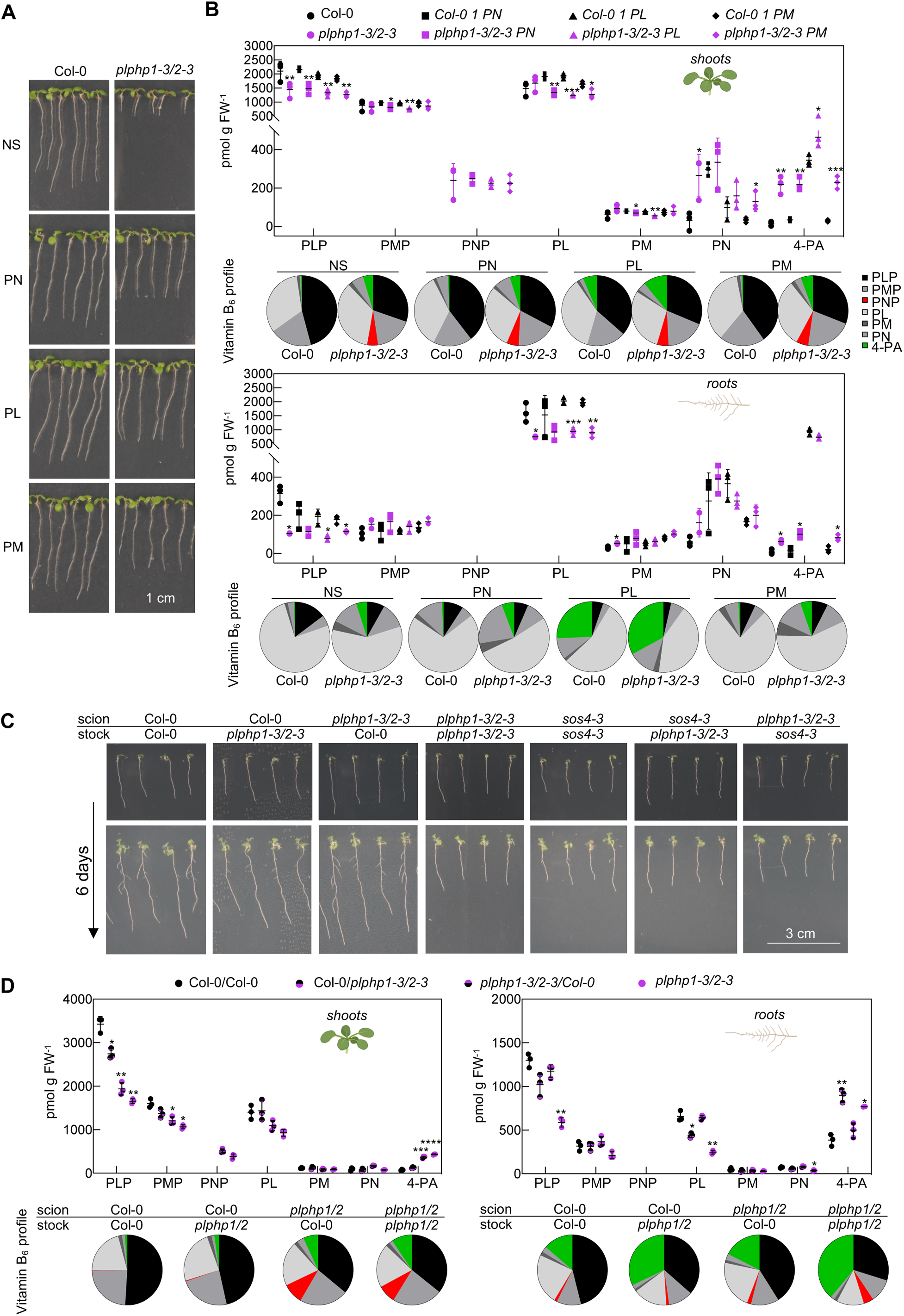
Vitamin B_6_ profiling of lines. **A)** Photos of 6 days old Col-0 (wild type) and *plphp1-3/2-3* mutant seedlings grown with full illumination on vertical culture plates under a 16-hour photoperiod (120 μmol photons m^−2^ s^−1^) at 22°C and 8 hours dark at 18°C without (NS) or with supplementation with either pyridoxine (PN), pyridoxal (PL) or pyridoxamine (PM) (5 μM in each case). **B)** Scatter dot plot (top) showing the abundance of seven vitamin B_6_ related compounds (pyridoxal 5ʹ-phosphate (PLP), pyridoxamine 5ʹ-phosphate (PMP), pyridoxine 5ʹ-phosphate (PNP), PL, PM, PN, 4-pyridoxic acid (4-PA)), in separated shoots and roots of 6 days old seedlings of lines as in **A**. Asterisks represent significant differences in the corresponding vitamer levels according to individual unpaired *t*-tests between genotypes within conditions. The same results are represented as pie charts on the bottom showing the relative distribution of the individual vitamers. **C)** Representative images of self-grafted and reciprocal grafts of seedlings of Col-0, *plphp1-3/2-3* or *sos4-3* as indicated right after grafting at 6 days after germination (top) and following a 6 days recovery period (bottom). Seedlings prior to grafting were grown with the root area shaded to provide comparable root morphology at the time of grafting and were illuminated after grafting. **D)** Vitamin B_6_ quantification in the shoots (left) and roots (right) of 28 days old self and reciprocal grafts between Col-0 and *plphp1-3/2-3* (3 weeks after grafting). Split circles represent reciprocal grafts with color and position indicating genotype and tissue, respectively: Col-0 (black), *plphp1-3/2-3* (purple), scion (top) and root stock (bottom). Letters represent significantly different groups (p≤0.05) according to an ordinary one-way ANOVA with Tukey’s multiple comparison test within vitamers. The same results are represented as pie charts on the bottom showing the relative distribution of the individual vitamers.

We were next prompted to investigate whether the presence of PLPHP and/or maintenance of the vitamin B_6_ profile of the shoot can influence root development. To test this notion, we performed reciprocal and self-grafting between Col-0 and *plphp1-3/2-3* that were first grown with the roots shaded to achieve equivalent development and then exposed to illumination following the grafting procedure. We observed that the wild type scion could restore the development of the *plphp1-3/2-3* mutant rootstock upon illumination (Figure 5C). Notably, the mutant scion did not disrupt the development of the wild type rootstock (Figure 5C), indicating that a shoot derived factor in *plphp1-3/2-3* does not impair its own root development under this condition. For example, the potential negative impact of the accumulated PNP in the shoot that could inhibit PLP-dependent enzymes (Tschopp and Kirschner, 1980; Ito et al., 2019) can be ruled out. Profiling of the B_6_ vitamer content of shoot and root tissue of the grafted plants illustrated that the genotype specific tissue features of intact or self-grafted plants remained similar in the reciprocal grafts with the striking exception of PLP (Figure 5D). Specifically, in the reciprocal grafts of wild type and *plphp1-3/2-3*, the level of PLP decreased in the grafted wild type tissue concomitant with an increase in the corresponding grafted tissue of *plphp1-3/2-3* (Figure 5D). By contrast, other vitamer changes diagnostic of *plphp1-3/2-3* shoot (increase in PNP) or root tissue (decrease of PL) remained similar even when grafted to wild type (Figure 5D). These observations suggested that the major cause of the *plphp1-3/2-3* root phenotype is related to PLP. In particular, it becomes clear that the presence of PLPHP itself in either tissue is sufficient to maintain PLP levels, with the root tissue being most sensitive to the deficit in PLP and/or PLPHP upon illumination. Remarkably, in the absence of PLPHP the deficit of PLP can be serviced distally and restores root growth in illuminated roots of *plphp1-3/2-3*. To test the notion of distal servicing of PLP to restore root growth in *plphp1-3/2-3* under illumination, we performed reciprocal grafts of *plphp1-3/2-3* and the PL/PN/PM kinase mutant *sos4-3* that suffers from a PLP-deficiency in shoots and roots (Gorelova et al., 2022) but has functional PLPHPs. The experiment was conducted as above by first growing seedlings with roots shaded and under low light during which *sos4-3* behaves like wild type (Gorelova et al., 2022) and then exposure of roots to illumination following the grafting procedure. Unlike wild type, scions of *sos4-3* could not rescue the root growth defect of *plphp1-3/2-3* (Figure 5C), supporting the notion of distal servicing of PLP in roots of *plphp1-3/2-3*. Of note, we checked if the AtPLPHP1 protein (for which we have an antibody) is mobile from the shoot to the root, but did not detect a change in protein levels in root tissues of *plphp1-3/2-3* stocks grafted with wild type scions (Supplementary Figure S6).

We infer that root photomorphogenesis is sensitive to a deficit in PLP when PLPHPs are absent, which can apparently be serviced distally (implying sensing) when sufficient levels of PLP are present in the shoot. On the other hand, PLPHP is sufficient to maintain PLP status in either shoot or root tissues indicating that PLPHP plays a protective role for the highly reactive PLP coenzyme.

### Evidence that PLPHP1/2 is required for dynamic functioning of PLP-dependent enzymes

We next probed why the root tissue was particularly sensitive to the PLP status under illumination in *plphp1-3/2-3*. PLP functions as a coenzyme in a plethora of reactions involving transamination, decarboxylation and racemization, making it pivotal for metabolic homeostasis. To assess if the absence of PLPHP poses a general pressure on PLP-dependent enzyme activities, we monitored sensitivity of a range of Arabidopsis mutants that are deficient in PLP (*pdx1.1*, *pdx1.3*, *sos4-3*, *pdx3-3*, *plphp1-3/2-3*) to D-cycloserine (DCS). The latter is taken up by plant cells and forms a stable aromatic product with PLP in the corresponding enzyme active sites blocking catalysis (Amorim Franco et al., 2017; de Chiara et al., 2020). Interestingly, despite similar levels of PLP deficit among the mutant lines used (Supplementary Figure S7), the *plphp1/2* mutant exhibited higher sensitivity to DCS compared to the other lines or wild type (Figure 6A and B). Importantly, the complementing line *PLPHP1/2c* behaved similar to wild type indicating that hypersensitivity was indeed associated with the absence of PLPHPs in the corresponding mutant lines (Figure 6A and B). This could imply that despite the overall deficiency in PLP at the tissue level in *plphp1-3/2-3*, more proteins have PLP bound (thus forming a dead-end complex with DCS making the mutant hypersensitive) and/or there is a specific role for PLPHP in local PLP homeostasis and dynamics.

**Figure 6.**
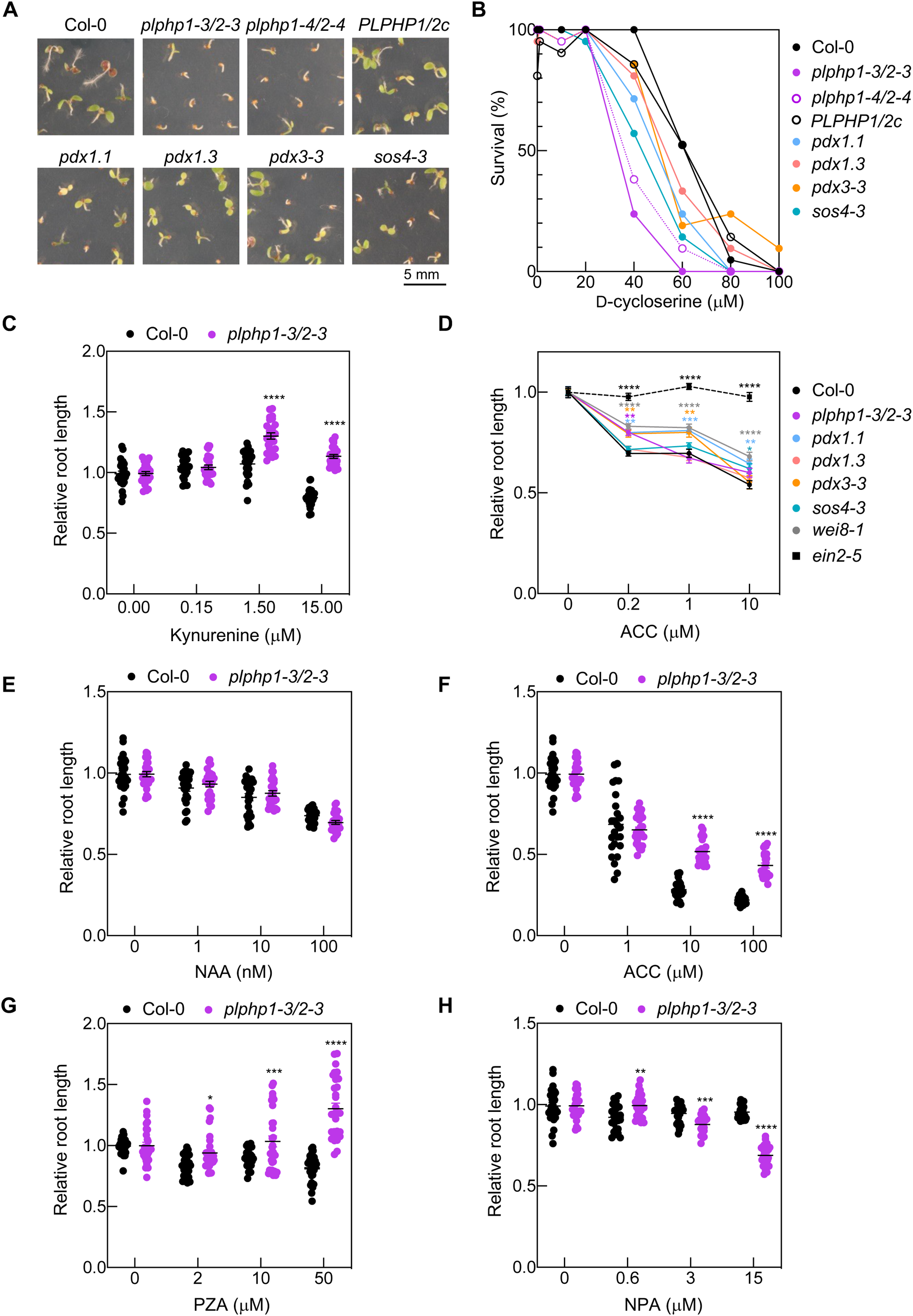
Treatments to assess the importance of PLP homeostasis by PLPHP. **A)** Representative images of the plant lines as indicated (Col-0 is the wild type; CRISPR mutants *plphp1-3 plphp2-3*, *plphp1-4 plphp2-4*; *plphp1-3/2-3* carrying both *PLPHP1* and *PLPHP2* (*PLPHP1/2c*) transgenes; *pdx1.1* and *pdx1.3* (biosynthesis *de novo* mutants); as well as *pdx3-3* and *sos4-3* (salvage pathway mutants) 9 days after germination on culture plates supplemented with 60 μM D-cycloserine (DCS) grown under a 16-hour photoperiod (120 μmol photons m^−2^ s^−1^) at 22°C and 8 hours dark at 18°C. **B)** Percentage survival of the indicated plant lines (as in **A**) on culture plates supplemented with increasing concentrations of DCS (n=21), grown as in **A** and 9 days after germination. **C)** Relative length of illuminated roots of the indicated lines (Col-0 is the wild type; CRISPR mutant *plphp1-3 plphp2-3*) grown on culture plates supplemented with the increasing concentrations of kynurenine compared to their root length without supplementation, grown as in **A** and 4 days after germination. Data represents the mean of 30 biological replicates ± SEM. **D)** Relative root length of etiolated seedlings of the indicated plant lines (as in **A**, as well as the local auxin biosynthesis mutant *wei8-1* and ethylene insensitive mutant *ein2-5*) grown on culture plates for 4 days and supplemented with increasing concentrations of 1-aminocyclopropane-1-carboxylic acid (ACC) relative to their root length without supplementation. Data represents the mean of 45 biological replicates ± SEM. **E-G)** Relative length of illuminated roots of grown on culture plates under a 16-hour photoperiod (120 μmol photons m^−2^ s^−1^) at 22°C and 8 hours dark at 18°C supplemented with increasing concentrations of either 1-napthaleneacetic acid (NAA, **E**), or 1-aminocyclopropane carboxylic acid (ACC, **F**), or pyrazinamide (PZA, **G**) or naphthylphthalamic acid (NPA, **H**), relative to their root length without supplementation. Data represents the mean of 30 biological replicates ± SEM. Statistical analyses were performed by multiple unpaired *t*-tests with Holm-Sidak’s correction for multiple comparison and asterisks mark significant difference from Col-0 (ns>0.05, *p≤0.05, **p≤0.005, *p≤0.0005, ****p≤0.0001).

The phytothormones ethylene and auxin are crucial for root tip growth and development and moreover, their homeostasis is particularly important for adaptive developmental decisions for example in the response to illumination (Růžička et al., 2007; Stepanova et al., 2007). In this context, it is significant that PLP is a coenzyme for several of the enzymes involved in the biosynthesis of these hormones, e.g. TRYPTOPHAN AMINOTRANSFERASE OF ARABIDOPSIS (TAA1) for auxin and 1-AMINOCYCLOPROPANE-1-CARBOXYLIC ACID (ACC) SYNTHASE for ethylene, respectively (Boller et al., 1979; Tao et al., 2008). Moreover, optimization of auxin and ethylene for varying environmental conditions is coordinated by REVERSAL OF *sav3* phenotype (VAS1), a protein that is also dependent on PLP (Zheng et al., 2013). Thus, while it would be difficult to dissect a specific response given the plethora of reactions PLP is involved in, we engaged a notion that PLPHPs may be pertinent under circumstances where a dynamic/adaptive requirement for PLP might be required, such as root illumination. This notion is not without precedent as recently enzyme activity of TAA1, involved in local auxin biosynthesis in the root meristematic zone, was shown to be dynamically regulated as a function of PLP (Wang et al., 2020). Specifically, TAA1 is alternately “PLPylated” or phosphorylated within the active site coincident with active and inactive states depending on the requirement for local auxin, although how PLP occupancy is controlled is not known. Local auxin biosynthesis has recently been reported to be inhibited upon root illumination (Spaninks and Offringa, 2023). In the first instance, we tested the response to the TAA1 inhibitor L-kynurenine (He et al., 2011) and observed that illuminated root growth was improved in *plphp1-3/2-3* in contrast to wild type (Figure 6C). Speculatively, this might suggest that PLPHP is required to sequester PLP from TAA1. On the other hand, local auxin biosynthesis is required for the so-called triple response to ethylene application in etiolated seedlings that includes shortening of the root (Stepanova et al., 2008). Upon application of the ethylene precursor ACC, with the exception of reduced sensitivity at the lowest concentration used, *plphp1-3/2-3* did not show any significant difference in the root length of etiolated seedlings compared to wild type (Figure 6D). This is in contrast to *ein2-5* and *wei8-1* which clearly displayed insensitivity and reduced sensitivity to ethylene, respectively, compared to wild type (Figure 6D), consistent with the literature (Guzmán and Ecker, 1990; Stepanova et al., 2008). Interestingly, mutants of PLP biosynthesis and salvage (*pdx1.1* and *pdx3-3*, in particular) were less sensitive to ACC inhibition of etiolated root growth (Figure 6D). While these observations suggest that PDX1.1 might play a role in providing PLP to TAA1, akin to the previously proposed role of PDX3 (Kim et al., 2018), they suggest that the role of AtPLPHP1/2 is more pertinent under root illumination, in line with the phenotypic observations.

Next to test perturbation of auxin/ethylene homeostasis in the illuminated root of *plphp1-3/2-3*, we gauged the response to alterations in levels of either hormone by application of the synthetic auxin, 1-napthaleneacetic acid (NAA) or the ethylene precursor ACC, over a range of concentrations. While illuminated root growth of *plphp1-3/2-3* was similar to wild type upon application of NAA (Figure 6E), the response to ACC was less sensitive (Figure 6F). Furthermore, while illuminated root growth of the wild type was compromised in the presence of increasing concentrations of the ACC oxidase inhibitor pyrazinamide (Sun et al., 2017), that of *plphp1-3/2-3* was considerably improved (Figure 6G). In addition, illumination of roots leads to production of flavonols that inhibit polar auxin transport influencing levels in the root (Silva-Navas et al., 2016). We observed that illuminated root growth of *plphp1-3/2-3* was more sensitive than wild type to treatment with the polar auxin transport inhibitor naphthylphthalamic acid (Galweiler et al., 1998) (Figure 6H). We infer that auxin and ethylene homeostasis is perturbed in root tips of *plphp1-3/2-3* under illumination.

Taken together, the hypersensitivity to DCS, complex response to treatments that interfere with auxin and ethylene homeostasis particularly under root illumination allow us to infer that PLPHP is required for the adaptive response under these conditions. Notably, our data suggest that the problem is not a deficit in PLP *per se* but rather regulation of homeostasis and PLP dynamics by PLPHP.

### Arabidopsis PLPHP can coordinate both PLP transfer and withdrawal

Monomeric PLPHPs with solvent exposed PLP as illustrated above, could be expected to service transfer to, or acceptance of, PLP from enzymes dependent on it to facilitate activation or deactivation, respectively. To investigate the possibility of PLP transfer, an *in vitro* assay was designed based on the activity of the PLP-dependent enzyme D-amino acid aminotransferase (D-AAT) (Ota et al., 2015). In the presence of D-Alanine and 2-oxoglutarate, the holoenzyme form of D-AAT (i.e. with PLP bound) can catalyze the formation of pyruvate and glutamate. Pyruvate formation is measured by addition of salicylaldehyde that leads to an orange chromophore that absorbs at 480 nm (Figure 7A). To facilitate the assay, an apo-enzyme form of D-AAT (apo-D-AAT) enzyme was prepared by treatment with DCS and buffer exchange (Figure 7B). In the first instance, we compared the relative activity of the holoenzyme form of D-AAT (holo-D-AAT) to the apoenzyme form and observed that the latter could be reactivated by addition of molar quantities of commercial PLP (Figure 7C). Significantly, addition of freshly purified AtPLPHP1 with PLP bound (holo-PLPHP) could also reactivate the apo-enzyme preparation of D-AAT (Figure 7D). By contrast, an apo-protein preparation of AtPLPHP1 (apo-PLPHP) failed to reconstitute activity to the apo-D-AAT (Figure 7D). This data suggests that holo-PLPHP can transfer PLP to enzymes dependent on the coenzyme for catalysis. To test the counter scenario, that is if AtPLPHP1 could accept PLP from a holoenzyme, apo-PLPHP was added to the reaction containing holo-D-AAT, which resulted in a decrease in the activity of the latter (Figure 7E). This indeed suggested that AtPLPHP1 is also capable of accepting PLP. Moreover, addition of the AtPLPHP1 K55A mutant, which cannot form the Schiff’s base with PLP, did not alter the activity of the reaction containing holo-D-AAT (Figure 7E).

**Figure 7.**
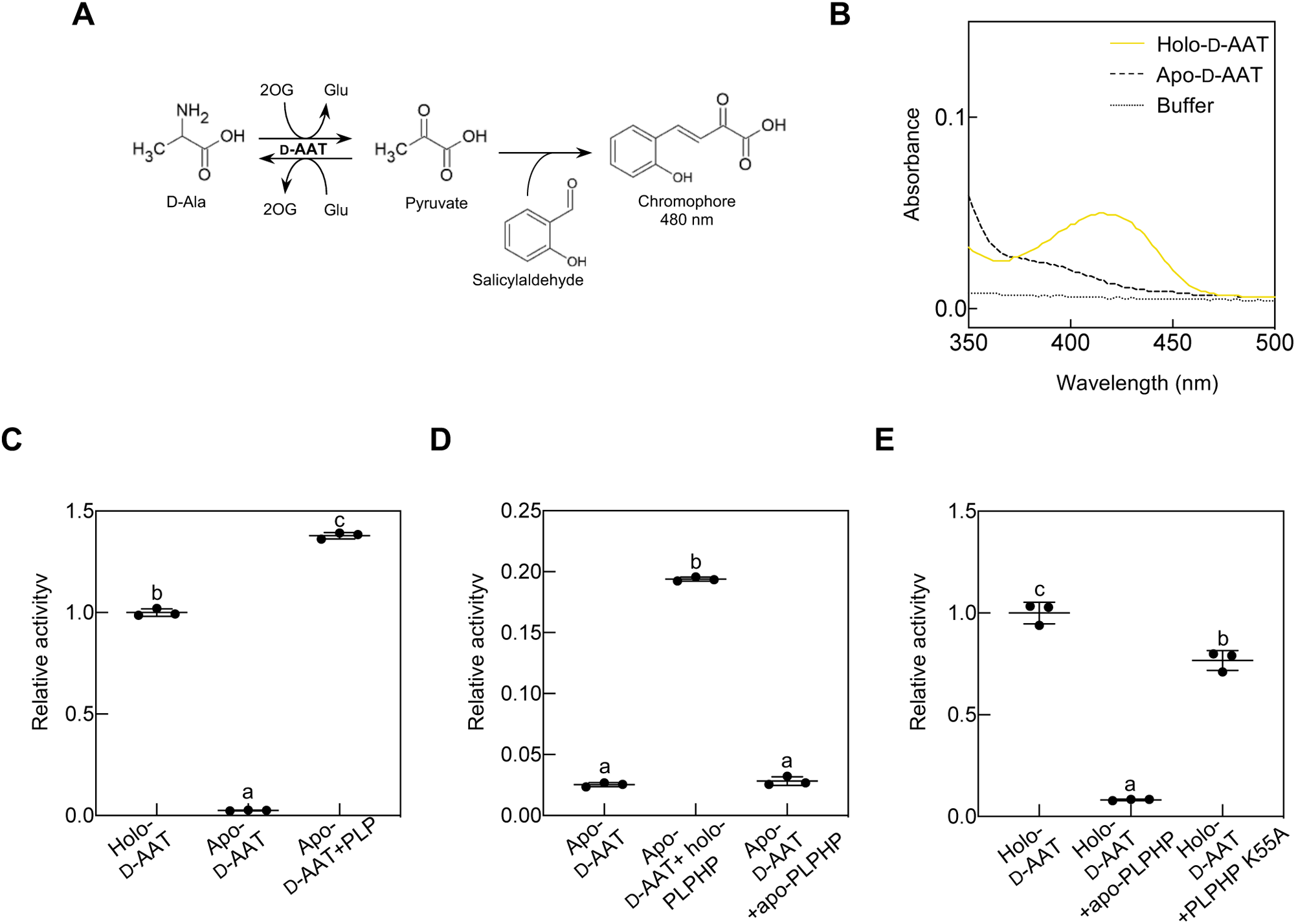
Enzyme assays to assess PLPHP functionality. **A)** Scheme of the activity assay of D-amino acid aminotransferase (D-AAT). In the presence of D-Alanine (D-Ala) and 2-oxoglutarate (2OG), D-AAT catalyzes the formation of pyruvate and glutamate (Glu). Pyruvate formation is measured by addition of salicylaldehyde that leads to an orange chromophore that absorbs at 480 nm. **B)** Absorption spectra of D-AAT as isolated (yellow; Holo-D-AAT), and after treatment with D-cycloserine and buffer exchange to remove PLP (black dashed line; Apo-D-AAT), compared to the buffer (black dotted line; 50 mM HEPES buffer at pH 7.4 containing 300 mM sodium chloride). **C)** Relative activity of Holo-D-AAT, or Apo-D-AAT without or with addition of PLP. **D)** Relative activity of Apo-D-AAT in the absence or presence of either AtPLPHP1 as purified (holo-PLPHP), or the apoenzyme form of AtPLPHP1 (apo-PLPHP) prepared as for apo-D-AAT. **E)** Relative activity of Holo-D-AAT in the absence or presence of either apo-PLPHP, or AtPLPHP1 with the PLP binding lysine residue mutated (PLPHP K55A). Values in **C**-**E** are represented as pyruvate produced relative to holo-D-AAT. Letters represent significantly different groups discovered by a Brown-Forsythe ANOVA test with Dunnett’s T3 multiple comparison (p≤0.05).

This data supports the notion that PLPHP has the capacity to both transfer and withdraw PLP suggesting a dynamic role within PLP homeostasis.

## DISCUSSION

PLPHP has been studied in a number of kingdoms most extensively in bacteria and mammalian cells but has been perceptibly ignored in plant systems. Here we provide insight into features of PLPHPs in *plantae*. In the first instance, we demonstrate that there are two homologs in higher plants, which is not reported for other kingdoms. Our studies show that at least in Arabidopsis, the two homologs appear redundant in terms of the observed morphological phenotype that is hypersensitivity of root growth to light. Although roots are typically shielded from light when grown in soil and only the upper soil layers permit limited light penetration, inferences can be deduced from the phenotypic response of illuminated roots. Similar to bacteria and mammals, perturbation of the levels of several B_6_ vitamers was observed demonstrating that the protein is required for homeostasis. While the vitamin B_6_ profiles are distinct for shoots versus roots both have a clear deficit in PLP. Remarkably, the PLP deficit in shoots does not manifest a morphological phenotype unlike disruption of the biosynthesis *de novo* or the salvage pathways. Moreover, micrografting studies suggest that the root PLP deficit can be serviced distally by the shoot. The hypersensitivity of roots to light in the absence of PLPHPs can be alleviated by manipulation of phytohormone levels (auxin and ethylene), strongly hinting that the proteins are required for achieving homeostasis under this condition. The ability of PLPHPs to accept or donate PLP, at least *in vitro*, while having no innate catalytic function, indicate a role in dynamic management of the coenzyme. Overall, our study indicates that PLPHPs contribute to surveillance of vitamin B_6_ homeostasis, likely acting as a rheostat in adaptive responses as a function of the use of the coenzyme PLP.

Our systematic analysis here, while revealing two homologs of PLPHP in higher plant species also reveals an interesting N-terminal extension feature in some PLPHP1 orthologs. This extension is not predicted to be an organellar targeting peptide according to several available algorithms (Predotar (Small et al., 2004), MitoFates (Fukasawa et al., 2015), TargetP2.0 (Almagro Armenteros et al., 2019) and SignalP5.0 (Almagro Armenteros et al., 2019)). Interestingly though, downstream of this extension a mitochondrial targeting sequence is predicted as well as at the N terminus of PLPHP2 (Supplementary Figure S8). Indeed, the mitochondrial targeting peptide is conserved in all Angiosperm PLPHPs, thus the upstream extension in certain PLPHP1 orthologs, may be a regulatory feature pertaining to subcellular localization that either requires processing or circumstantially prevents mitochondrial targeting. Notably, this extension is only found in certain eudicots whereas monocots lack the extension. In line with this, a recent study on a PLPHP1 homolog in rice reported mitochondrial localization (Cao et al., 2022). Indeed, PLPHP homologs from yeast and humans are reported to be localized to the mitochondria (Johnstone et al., 2019). Nonetheless, we show here that Arabidopsis PLPHP1 alone, which harbors the N-terminal extension, complements the morphological root phenotype of the double *plphp1/2* mutants indicating that it is functional *in vivo*. Further studies on this extension may unveil its precise role. On the other hand, PLPHP2 can also rescue the morphological phenotype observed upon root illumination in Arabidopsis. As gene homologs frequently act redundantly, it was only with the observation of two homologs in higher plants and disruption of both in Arabidopsis that we could observe the conditional phenotype reported here. Therefore, we assume redundancy at least in terms of hypersensitivity to root illumination in Arabidopsis. However, it should be noted that while rice has two homologs of PLPHP (Os01g72205 and Os05g05740) as elucidated here, the latter one was recently reported to confer heat tolerance at the heading stage of the wild rice variety *Oryza rufipogon* Griff. and thus named HTH5 (Cao et al., 2022). Analysis of the second rice homolog will reveal the level of redundancy between the paralogs.

Our study (summarized in Figure 8) clearly implicates PLPHPs in regulation of B_6_ vitamer levels in Arabidopsis. As PLPHP has no known enzyme activity, the deficit in PLP/PL concomitant with the increase in the degradation product 4-PA suggests a protective function perhaps to ensure supply of PLP or management of the coenzyme. This statement is supported by the inability to restore vitamer homeostasis upon feeding of the non-phosphorylated vitamers, even though these can be converted to PLP by salvage pathway enzymes. Dissection of responses in shoots versus roots of Arabidopsis in the absence of PLPHPs allowed us to diagnose causal morphological effects. In particular, while we saw decreased PLP levels in both shoots and roots, the increase in PNP was confined to shoots. However, neither the deficit in PLP nor the accumulation of PNP in shoots seemed to impact the plants at least under our standard growth conditions. This is in contrast to what is hypothesized in other organisms where PNP accumulation was proposed to be causal to phenotypes observed (Plecko et al., 2017; Ito et al., 2019; Vu and Downs, 2021); although this does not rule out that excessive PNP may relate to phenotypes not noted here with Arabidopsis. On the other hand, a deficiency in the PLP pool could cause wide-ranging effects. The activities of PLP-dependent proteins would be expected to be altered, the order of which would depend on the affinity for PLP, the saturation level at the time of the deficit and the turnover rate of the protein itself. These features would be difficult to dissect *in vivo* in a physiological context. Nonetheless, enzymes with low affinity for PLP and high turnover, would be expected to be most sensitive to PLP loss. Further, enzymes that have a dynamic requirement for PLP (e.g. TAA1 (Wang et al., 2020)) would be highly sensitive. Here root hypersensitivity to illumination in the absence of PLPHPs indicate that this is a condition where management of PLP by these proteins is required. Root illumination necessitates reconfiguration of key phytohormones auxin/ethylene to achieve homeostasis and optimal root growth under this condition (Růžička et al., 2007; Stepanova et al., 2007). The inability to reach an equilibrium is manifested in developmental defects, particularly in the root tip, as observed here with *plphp1/2* mutants. Indeed, alleviation of the phenotype by manipulation of auxin or ethylene biosynthesis (e.g. with L-kynurenine and pyrazinamide, respectively (He et al., 2011; Sun et al., 2017)), and hyposensitivity to the ethylene precursor under illumination suggest that these plants cannot reach homeostasis of these hormones. Thus, a physiological function can be deduced for PLPHPs in plants. Furthermore, as the roots of certain plants such as epiphytes or germinating seeds and those cultivated in hydroponic or aeroponic systems may grow under illumination, the finding has applied relevance. An interesting observation here is that PLP homeostasis in the root can be serviced by the shoot.

**Figure 8.**
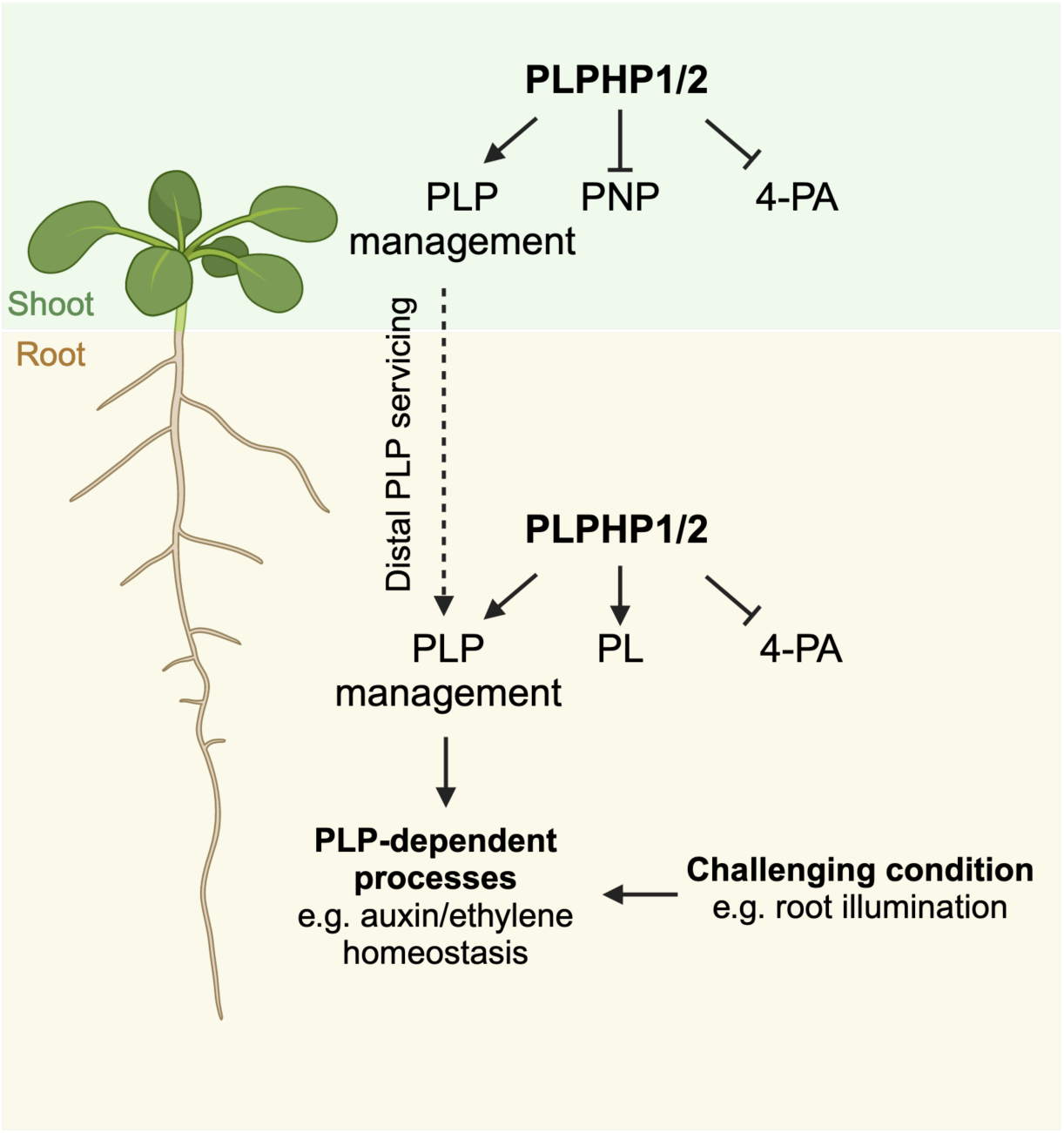
Scheme of findings in Arabidopsis. Two PLPHP homologs (PLPHP1/2) function in vitamin B_6_ homeostasis in Arabidopsis, by supporting pyridoxal 5’-phosphate (PLP) management in the shoot and the root, preventing the accumulation of pyridoxine 5’-phosphate (PNP) in the shoot and 4-pyridoxine acid (4-PA) in both tissues, as well as maintaining pyridoxal (PL) levels in the root. PLP management in the root serviced either by the local presence of PLPHP1/2 or by distal PLP servicing from the shoot is necessary under challenging conditions, exemplified by root illumination, which involves the maintenance of adaptive responses such as the homeostasis of auxin and ethylene. Created with Biorender.

This requires the presence of at least either PLPHP1/2 or SOS4 in the shoot, because in their absence root PLP homeostasis is not achieved. It remains to be determined if these proteins function alone or together and can detect distal aberrations in PLP levels. Thus, biosynthesis *de novo* is not sufficient for PLP supply in Arabidopsis plants and emphasizes the importance of PLP homeostasis for growth and development as serviced by salvage and homeostasis proteins. We postulate that management of PLP is not simply servicing supply or even a deficit but also implicates dynamic control of PLP occupancy of enzyme active sites. This statement is supported by the already mentioned elegant study of the local auxin biosynthesis protein TAA1 in which a switch for active versus inactive enzyme is operated through occupancy of the active site either by PLP (active) or P_i_ (inactive), where the latter occludes binding of the coenzyme (Wang et al., 2020). Interestingly, auxin itself mediates the phosphorylation via the kinase TMK4 forming a negative feedback loop (Wang et al., 2020). Reactivation of the TAA1 enzyme would require transfer of PLP. It is plausible that dynamic operation of this system to be coincident with local auxin requirements could be achieved by a protein such as PLPHP, which as we have shown has the ability to accept or donate PLP. Interestingly, dynamic phosphorylation and PLPylation is evolutionarily conserved across land plant species, and extends to ancestral species such as *Marchantia polymorpha* (Wang et al 2020). Although the only example so far, we hypothesize that many enzymes likely operate in this dynamic manner as such a system (i.e. plasticity in coenzyme use) bypasses the energy consuming requirement of holoenzyme replacement. Further exploration of such switches and the association of the response to environmental cues deserves attention.

In summary, we conclude that PLPHPs are important regulators of vitamin B_6_ homeostasis (likely in tandem with *de novo* and salvage pathways), which in turn is required for plant homeostasis to optimize growth and development coincident with environmental cues. We postulate that coordination of homeostasis is achieved as a function of coenzyme modulation, which operates in a dynamic equilibrium in a nexus of proteins that can control active site occupancy. This opens up a whole area of future investigation where the dynamic provision of a coenzyme could be seen as an adaptive response to an environmental challenge.

## MATERIAL AND METHODS

### Phylogenetic analysis

A list of 4886 amino acid sequences were retrieved from UniProt identified by K06997 orthology at the KEGG Orthology database (https://www.genome.jp/kegg/ko.html) (Kanehisa et al., 2016). The sequence list of Archaeplastida was refined by selecting representative species from 19 orders of Angiosperms and representatives of non-flowering clades, resulting in 4838 sequences including 79 from plants. A multiple sequence alignment was performed on the corresponding amino acid sequences using MAFFT v7.450 (Katoh and Standley, 2013) with BLOSUM62 scoring matrix. The phylogenetic tree was constructed with the Neighbor-Joining tree build method (Saitou and Nei, 1987) using the Jukes-Cantor genetic distance model with no outgroup. The multiple sequence alignment and tree construction was performed in Geneious® 11.1.5.

### Recombinant protein expression, purification and biochemical analyses

The full-length coding sequences of At*PLPHP1* (At1g11930) and At*PLPHP2* (At4g26860) were synthesized with codon optimization for expression in *E. coli* (Genscript) and cloned into the *pET-22b* expression vector (Novagen) for expression with a C terminal hexahistidine tag (*pET-AtPLPHP1* and *pET-AtPLPHP2*, respectively). Site directed mutagenesis (to give *pET-AtPLPHP1 K55A*) was performed using PfuTurbo AD high fidelity DNA polymerase (Agilent) and oligonucleotides as indicated in Supplementary Table S4. Constructs were transformed separately into the *E. coli* BL21 (DE3) strain and cultured in LB broth medium supplemented with 100 μg ml^−1^ ampicillin at 37°C. When the bacterial cultures reached an optical density of 0.5 at 600 nm, protein expression was induced by adding 100-500 μM of isopropyl β-D-1-thiogalactopyranoside followed by incubation with shaking for 3 hours at 37°C. The bacterial pellets were harvested by centrifugation and resuspended in lysis buffer (50 mM NaH_2_PO_4_ pH 8.0, containing 300 mM sodium chloride, 10 mM imidazole, and 0.1 mM phenylmethylsulfonyl fluoride and protease inhibitor cocktail (Roche)) and lysed by adding lysozyme followed by sonication. The soluble protein was first purified by affinity chromatography with Protino nickel nitrilotriacetic acid agarose (Macherey-Nagel) using the lysis buffer but with the imidazole concentration changed to sequential rounds of 20 mM and 250 mM for washing and eluting the protein, respectively. Additionally, the proteins were subjected to size exclusion chromatography on a Superdex 200 10/300 increase column (GE Healthcare) and eluted using 150 mM Tris–HCl pH 8.8 containing 300 mM sodium chloride. The *pKK223-3-Bsp.DAAT* (Ota et al., 2015) was a gift from Kazuaki Yoshimune (Nihon University, Japan) and the *pET-16(b)-BsPdxK* (Park et al., 2004) was a gift from Tadhg Begley (Texas A&M University, U.S.A.) with expression and purification as above. Purified recombinant proteins were buffer exchanged into either 50 mM HEPES pH 7.4 containing 300 mM sodium chloride or 50 mM sodium phosphate buffer pH 8.0 containing 300 mM sodium chloride using Amicon Ultra-15 centrifugal filter devices with a 10 kDa molecular weight cut-off.

### Spectrophotometric analyses

Absorption spectra were recorded in a Synergy 2 microplate reader (BioTek) using UV-Star flat bottom 96 well microplates (Grenier Bio-One) and 100 μl of 75 μM recombinant protein. Sodium hydroxide (0.1 M) was added directly to the protein sample. Sodium borohydride (0.5 mg ml^−1^) was added to an ammonium sulfate suspension of the protein followed by centrifugation at 15000 *g* for 15 min at 4°C and resuspended in 100 μl of 50 mM HEPES pH 7.4 containing 300 mM sodium chloride. Apoenzyme preparations of D-AAT and AtPLPHP1 were achieved by incubating 100 μM of purified protein with 75 mM D-cycloserine (Merck) in 50 mM HEPES pH 7.4 containing 300 mM sodium chloride on ice for 2 hours followed by buffer exchange as described above. The D-AAT assay was performed as outlined in (Ota et al., 2015) with slight modifications. The reaction mixture (150 μl total volume) consisted of 50 mM Tricine pH 8.0, containing 10 mM 2-oxoglutarate, 100 mM D-alanine, 1 μM D-AAT and (if applicable) 4 μM PLP (free or protein bound). After pre-incubation (1 min) the reaction was started by addition of 30 μl of 500 mM D-alanine and incubated for 10 min at 37°C. The reaction was stopped by the addition of 150 μl of 60% (w/v) potassium hydroxide and 50 μl 2% (w/v) salicylaldehyde in ethanol and incubation at 37°C for 10 min after which the absorbance at 480 nm was measured.

### Plant material and growth conditions

Arabidopsis (*Arabidopsis thaliana*) Col-0 ecotype was used from an in house stock and was the genetic background for all mutants used. Seeds of the T-DNA insertion lines *wei8-1* (N16407; At1g70560 (Stepanova et al., 2008)), *sos4-3* (N485452; At5g37850 (Gorelova et al., 2022)), *pdx1.1-1* (SM_3_22656; At2g38230; (Tambasco-Studart et al., 2005)), *pdx1.3* (N586418; At5g01410 (Tambasco-Studart et al., 2005)), *pdx3-3* (N659883; At5g49970 (Colinas et al., 2016)), *plphp1-1* (N829537; At1g11930) and *plphp2-1* (N470954; At4g26860) were obtained from in house stocks or the Nottingham Arabidopsis Stock Centre in which case plants homozygous for the T-DNA insertion were isolated (see Supplementary Table S4 for oligonucleotides used). Additional mutant alleles for *AtPLPHP1/2* were generated for both loci using a modified CRISPR/Cas9 protocol (Ursache et al., 2021), as well as complementing lines (see below). On soil, plants were either grown in pots or rhizoboxes on Einheitserde, Patzer classic ton Kokos that has a composition of 25% clay, 45% wheat peat, 15% brown peat, and 15% coconut fiber pH 5.8. Before use, the soil was sterilized for 1 hour at 60°C and supplemented once with a 2 ml l^−1^ solution of entomopathogenic nematodes (Andermatt, Traunem) and 2 tablespoons l^−1^ of *Bacillus thuringiensis israelensis* (Andermatt, Solbac). For growth on sterile culture, half-strength Murashige and Skoog medium (Murashige and Skoog, 1962) supplemented with 0.1 g l^−1^ myo-inositol and 0.5 g l^−1^ 2-(N-morpholino)ethanesulfonic acid (MES) at pH 5.7 and 8 g l^−1^ plant agar (Duchefa), autoclaved at 120°C and 2 bar for 20 minutes was used. Supplementation of media with the non-phosphorylated B_6_ vitamers PL/PN/PM (Merck) was performed by addition of the compounds at a concentration of 5 μM to the sterile culture plates. 1-naphthalene acetic acid (Duchefa), 1-aminocyclopropane (Merck), L-kynurenine (Merck), pyrazinamide (Merck) and naphthylphthalamic acid (Merck) were supplemented at concentrations indicated in the corresponding figures from 10^3^ concentrated stocks dissolved in dimethylsulfoxide (Acros). Supplementation with D-cycloserine (Merck) was performed on horizontal 6 cm diameter petri dishes with half strength MS medium supplemented with the concentration of D-cycloserine indicated in the figures. In all cases, plants were grown under a 16-hour photoperiod at approximately 120 μmol photons m^−2^ s^−1^ generated by fluorescent lamps at 22°C and 8-hour darkness at 18°C with 60-70% relative humidity and ambient CO_2_ generally in Percival growth chambers (CLF Plant Climatics GmbH) or in house walk-in rooms. In each case, seeds were surface sterilized by washing with 70% (v/v) ethanol and were stratified for 3 days at 4°C in the dark before transferring to photoperiodic conditions. Plant material was harvested 4 hours after the onset of illumination for all analyses.

### Generation of mutant alleles by CRISPR/Cas9 technology

Three target sites were selected for each loci (At1g11930, At4g26860) using a corresponding plugin in Geneious® 11.1.5, prioritizing candidate sites with the highest activity score (Doench et al., 2014) and specificity score (Ran et al., 2013) mapping against the reference genome of Arabidopsis. Briefly, protospacers (see Supplementary Table S4) were annealed into Golden-Gate entry vectors (*pGG-A-AtU6-26-BbsI-ccdB-BbsI-B, pGG-B-AtU6-26-BbsI-ccdB-BbsI-C, pGG-C-AtU6-26-BbsI-ccdB-BbsI-D, pGG-D-AtU6-26-BbsI-ccdB-BbsI-E, pGG-E-AtU6-26-BbsI-ccdB-BbsI-F, pGG-F-AtU6-26-BbsI-ccdB-BbsI-G*) (Decaestecker et al., 2019) using the BbsI Golden Gate reaction creating sgRNAs under the expression of the Arabidopsis U6 promoter. These cassettes were assembled in the vector *pDONR221-attL1-A-ccdB-G-attL2* (generated here) using the BsaI Golden Gate reaction and finally recombined by an LR Gateway reaction into the T-DNA region of the RU051 vector that contains *Streptococcus pyogenes* Cas9 driven by the *Petroselinum crispum* (parsley) *UBI4-2* promoter as well as a red fluorescent protein (FastRed), expression of which is driven by the seed specific promoter of *OLEOSIN1* (*OLE1*, At4g25140) as a plant selection marker (Ursache et al., 2021). The respective constructs were transformed into Arabidopsis using the floral dip method (Clough and Bent, 1998) and *Agrobacterium tumefaciens* strain GV3101 (*C58C1* (*RifR*)*; pMP90* (*pTiC58ΔT-DNA*) (*GentR*)). Screening of plants homozygous for a mutation in either *AtPLPHP1*, *AtPLPHP2* or both genes was performed by extracting gDNA from rosette leaves of individual plants grown from red fluorescent T1 seeds, and PCR using oligonucleotides amplifying a region spanning across the three target sites per gene (Supplementary Table S4). Plants carrying mutations in the target genes were selected and T2 seeds negative for red fluorescence indicative of loss of the transgene were chosen and validated by PCR. Specifically, a line carrying a genetic reversion between nucleotide residues 106-261 and a 951 bp deletion between residues 261-1057 in *AtPLPHP1* was annotated as *plphp1-2*; a line with a 1573 bp deletion from 12 bp upstream of the start codon until 158 bp downstream of the stop codon in *AtPLPHP2* removing the entire coding sequence was annotated as *plphp2-2*; two lines with deletions in both *AtPLPHP1* and *2* were isolated, *plphp1-3/2-3* with a 155 bp deletion between 106-261 in *AtPLPHP1* and 335 bp deletion between 35-370 in *AtPLPHP2* and *plphp1-4/2-4* with a 312 bp deletion and a 8 bp insertion between 261-573 in *AtPLPHP1* and a 228 bp deletion between 2-236 in *AtPLPHP2*, respectively (Figure 3A).

### Construction of other transgenic lines

For complementation with either *AtPLPHP1*, *AtPLPHP2* or both homologs, the genes were amplified from Arabidopsis genomic DNA (Col-0) (3363 bp from locus At1g11930 (*AtPLPHP1*): from 1533 bp upstream to 1830 bp downstream of the start codon and 2346 bp from locus At4g26860 (*AtPLPHP2*): from 683 bp upstream to 1660 bp downstream of the start codon), and inserted into the T-DNA region of binary vector *pFASTR-A-G* carrying the FastRed selection marker under the expression of the seed specific *OLEOSIN1* promoter (Decaestecker et al., 2019). The constructs were transformed into the double mutant *plphp1-3/2-3* Arabidopsis using the floral dip method (Clough and Bent, 1998) and *Agrobacterium tumefaciens* strain GV3101 and the resulting transformants were segregated for homozygous single T-DNA insertion by analyzing the segregation of the red fluorescent seeds in the three successive generations following transformation.

### Gene expression analyses

RNA was extracted from 6-days-old seedlings grown in sterile culture using the RNA NucleoSpin Plant kit (Macherey-Nagel) according to the manufacturer’s instructions with on-column DNase treatment. Reverse transcription (RT) was performed using 1 μg of total RNA as template and Superscript II (Invitrogen) according to the instructions: stock oligo(dT)_15_ primer (C1101, Promega) concentration was 100 ng/μl, and 0.5-μl Superscript II enzyme was used per reaction. RT-qPCR was performed in 384-well plates on a 7900HT Fast Real-Time PCR system (Applied Biosystems) using PowerUp SYBR Green master mix (Applied Biosystems), 1 ng cDNA as template and the following amplification program: 10-min denaturation at 95°C followed by 40 cycles of 95°C for 15 s and 60°C for 1 min. The data was analyzed using the comparative cycle threshold method (2^−ΔΔCT^) normalized to the reference gene *PP2AA3* (At1g13320). Oligonucleotide pairs used are indicated in Supplementary Table S4. For immunochemical analyses, protein was extracted from frozen ground plant tissues in 50 mM sodium phosphate pH 7.0 containing 10 mM EDTA, 0.1% (v/v) Triton X-100, 5 mM β-mercaptoethanol and 1% (v/v) complete protease inhibitor cocktail (Merck). The protein concentration was quantified in crude extracts with the Bradford assay (Bradford, 1976) and equal amounts of protein were boiled for 4 min in sample buffer (50 mM Tris-HCl pH 6.8 containing 1% SDS (w/v), 3% glycerol (v/v), 0.01% bromophenol blue (w/v) and 143 mM β-mercaptoethanol) and separated on a 13% SDS-PAGE. Proteins were then transferred onto a nitrocellulose membrane using the iBlot® Dry Blotting System (Life Technologies) and stained with Ponceau S (0.1% Ponceau S (w/v) in 5% acetic acid (v/v)) before blocking with 5% (w/v) low-fat milk in 25 mM Tris-HCl pH 7.5 containing 50 mM NaCl, 2.5 mM KCl, 0.05% (v/v) Tween 20 (TBST) for 1.5 hours. A polyclonal antibody was raised against the recombinant AtPLPHP1 in rabbits by BioGenes GmbH, and was purified against the recombinant protein and used in a 1:1000 dilution in the blocking solution in an overnight incubation at 4°C. The membrane was washed in TBST four times and incubated with goat anti-rabbit secondary antibody (Agrisera) conjugated to horseradish peroxidase in a 1:3000 dilution in blocking solution for 1 hour at room temperature (ca. 25°C). α-ACTIN-2 at a 1:10000 dilution was used as a loading control (Sigma-Aldrich) and the secondary antibody goat Anti-Mouse IgG (H + L)-HRP conjugate (Bio-Rad) was used at a dilution of 1:5000. The bands were visualized with Western Bright ECL (Advansta) on an Amersham Imager 680 (GE Life Sciences).

### Root length experiments

For root length measurements, seedlings were grown on vertical plates in a single horizontal line and the development of the root was recorded daily using either a Leica MZ16 stereomicroscope equipped with an Infinity 2 camera or a Canon EOS 77D camera fitted to a copy-stand. The length of the roots was measured with ImageJ https://imagej.net/ij/. Root shading was achieved by inserting the plates in a cardboard box lined with black duct tape, covering the plate area below the line of the seeds.

### Micrografting of Arabidopsis

Reciprocal and self-grafting was performed on 6-days-old Arabidopsis seedlings that were grown vertically on culture plates with the roots shaded. The hypocotyls were cut with a Wilkinson blade ∼1/3 distance from the cotyledons with respect to the length of the hypocotyl and perpendicular to the axis of the hypocotyl, after which one cotyledon was removed and the scions were secured on the corresponding rootstocks using sterile plastic tubing (ca. 1 mm). The grafts were transferred to recovery conditions (16-hour photoperiod at approximately 120 μmol photons m^−2^ s^−1^ generated by fluorescent lamps at 27°C and 8-hour darkness at 23°C with 60% relative humidity) with roots exposed to light and the root development was recorded for 9 days. After 10 days, successful grafts (with no adventitious root formation) were transferred to rhizoboxes (Gorelova et al., 2022) and grown for a further 7 days, roots and shoots were harvested separately in three biological replicates, each consisting of material from at least three plants to facilitate vitamin B_6_ profiling.

### Vitamin B_6_ analysis by HPLC

Vitamin B_6_ analysis was performed as described in (Steensma et al., 2023). For production of pyridoxine 5′-phosphate, which is not commercially available, a reaction mixture consisting of 50 mM Tris-HCl pH 8.0 containing 2 mM MgCl_2_, 2 mM ATP, 1 mM pyridoxine and 4 μM BsPdxK (Park et al., 2004) in a total volume of 500 μl was incubated overnight at 37°C, filtered with an Amicon Ultra-0.5 centrifugal filter device with 3 kDa molecular weight cut-off and analyzed by HPLC to verify completion of the reaction and was used as a standard.

### Analysis software

Data rendering and statistical analysis were performed using GraphPad Prism version 8.2.0 for Windows, GraphPad Software, San Diego, California USA, www.graphpad.com. Root length and cotyledon area quantification were performed by ImageJ (Schindelin et al., 2012) https://imagej.nih.gov/ij/. Multiple sequence alignments and phylogenetic analyses were performed in Geneious version 11.1.5. 3D protein structures were generated using Pymol version 4.6 (https://sourceforge.net/projects/pymol/).

### Accession numbers

Sequences of genes involved in this study can be found at The Arabidopsis Information Resource (https://www.arabidopsis.org/) under the following accession numbers: *EIN2* (At5g03280), *PDX1.1* (At2g38230), *PDX1.3* (At5g01410), *PDX3* (At5g49970), *PLPHP1* (At1g11930), *PLPHP2* (At4g26860), *SOS4* (At5g37850), *TAA1*/*WEI8* (At1g70560).

## Acknowledgments

We thank Benoît Maillot (University of Geneva) for his assistance with aspects of recombinant protein expression and purification, and Sylvain Loubéry (University of Geneva) for help with microscopy. We are indebted to Mireille de Meyer Fague (University of Geneva) for assistance with plant methods. We gratefully acknowledge Kazuaki Yoshimune (Nihon University, Japan) for the gift of *pKK223-3-Bsp.DAAT*, Marie Barberon (University of Geneva) for the RU051 vector and Tadhg Begley (Texas A&M University, U.S.A.) for the *pET-16(b)-BsPdxK*.

## Author contributions

P.F. and T.B.F. carried out experimental work, contributed tools, and analyzed data. T.B.F. secured funding, supervised the research and wrote the paper.

---

**Supplementary Figure S1.**
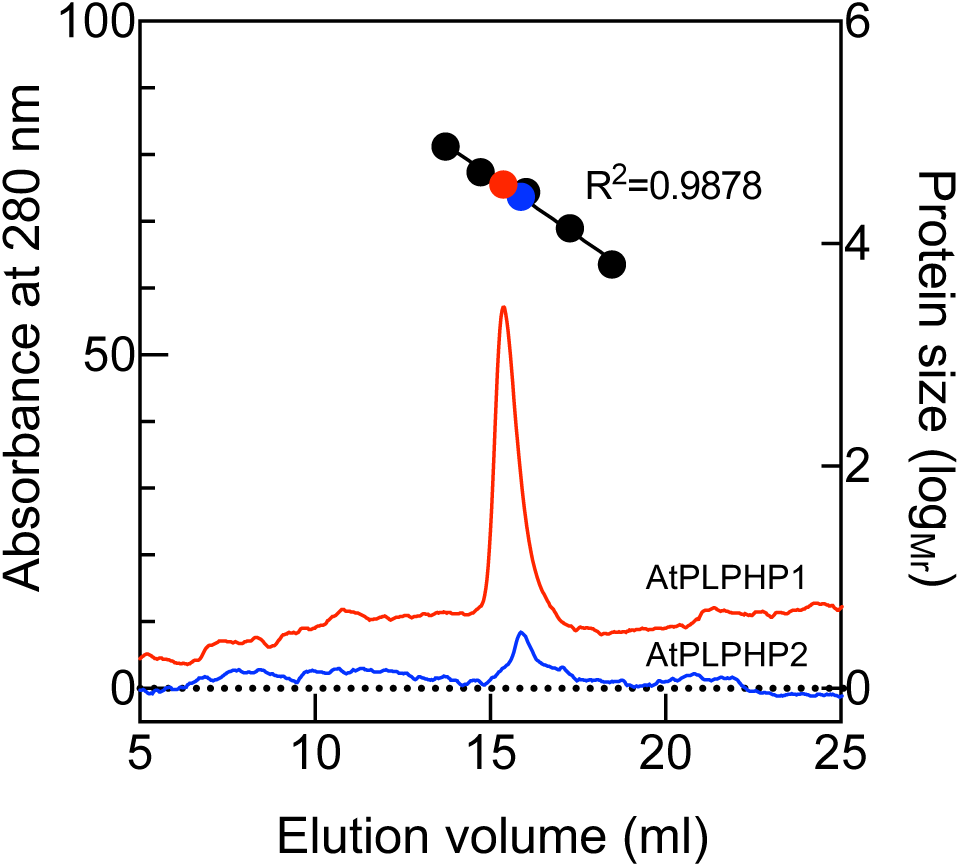
AtPLPHPs behave as monomers. Size exclusion chromatography of recombinant AtPLPHP1 (red) and AtPLPHP2 (blue). The apparent molecular mass of the proteins (shown as log molecular mass) was calculated based on the elution volume of molecular mass standards (black dots) consisting of aprotinin (6500 Da), ribonuclease A (13700 Da), carbonic anhydrase (29000 Da), ovalbumin (43000 Da) and conalbumin (75000 Da), while blue dextran 2000 (2×10^6^ Da) was used as an indicator of void volume (black dashed line) as provided in a Gel filtration LMW calibration kit (GE Healthcare).

**Supplementary Figure S2.**
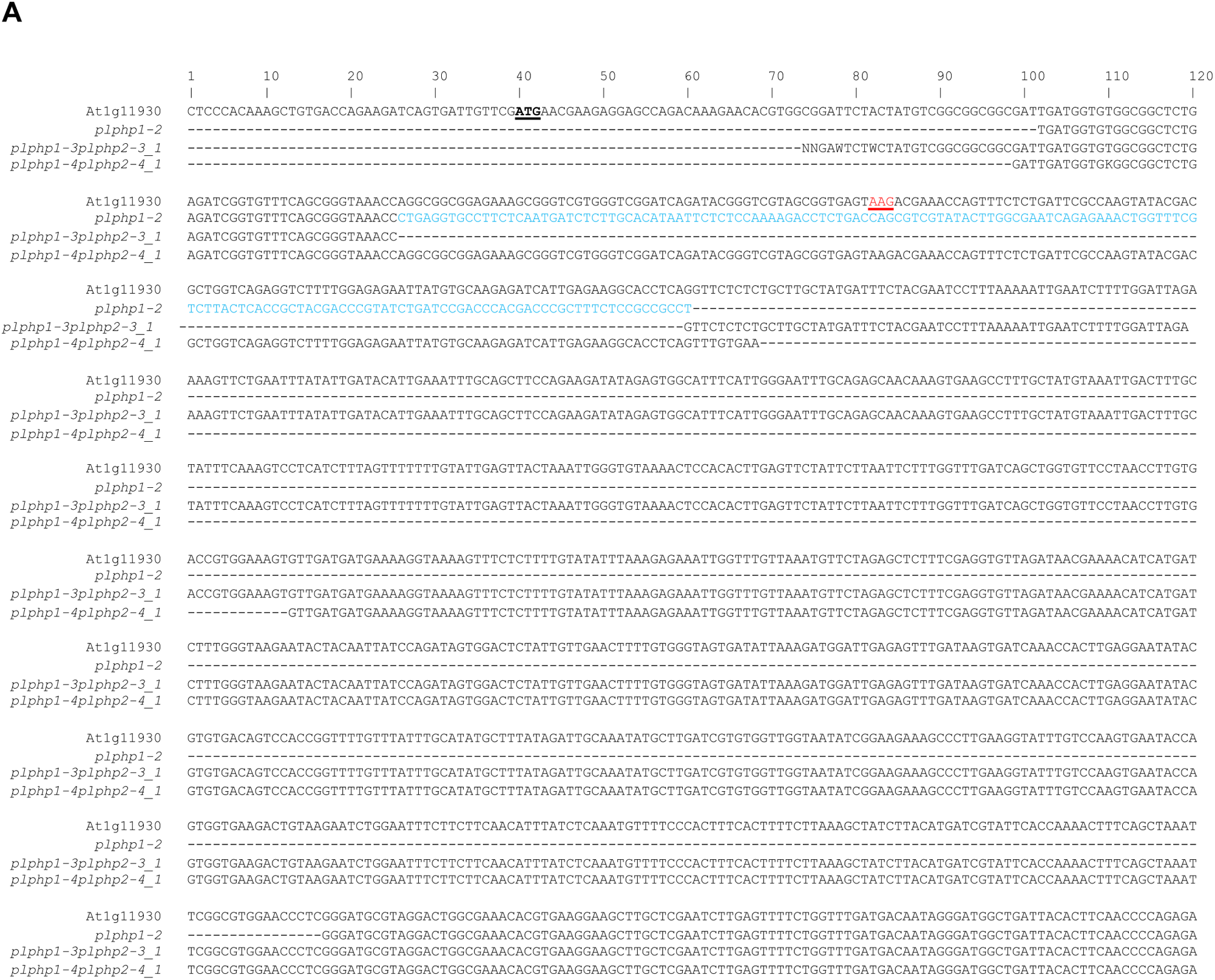
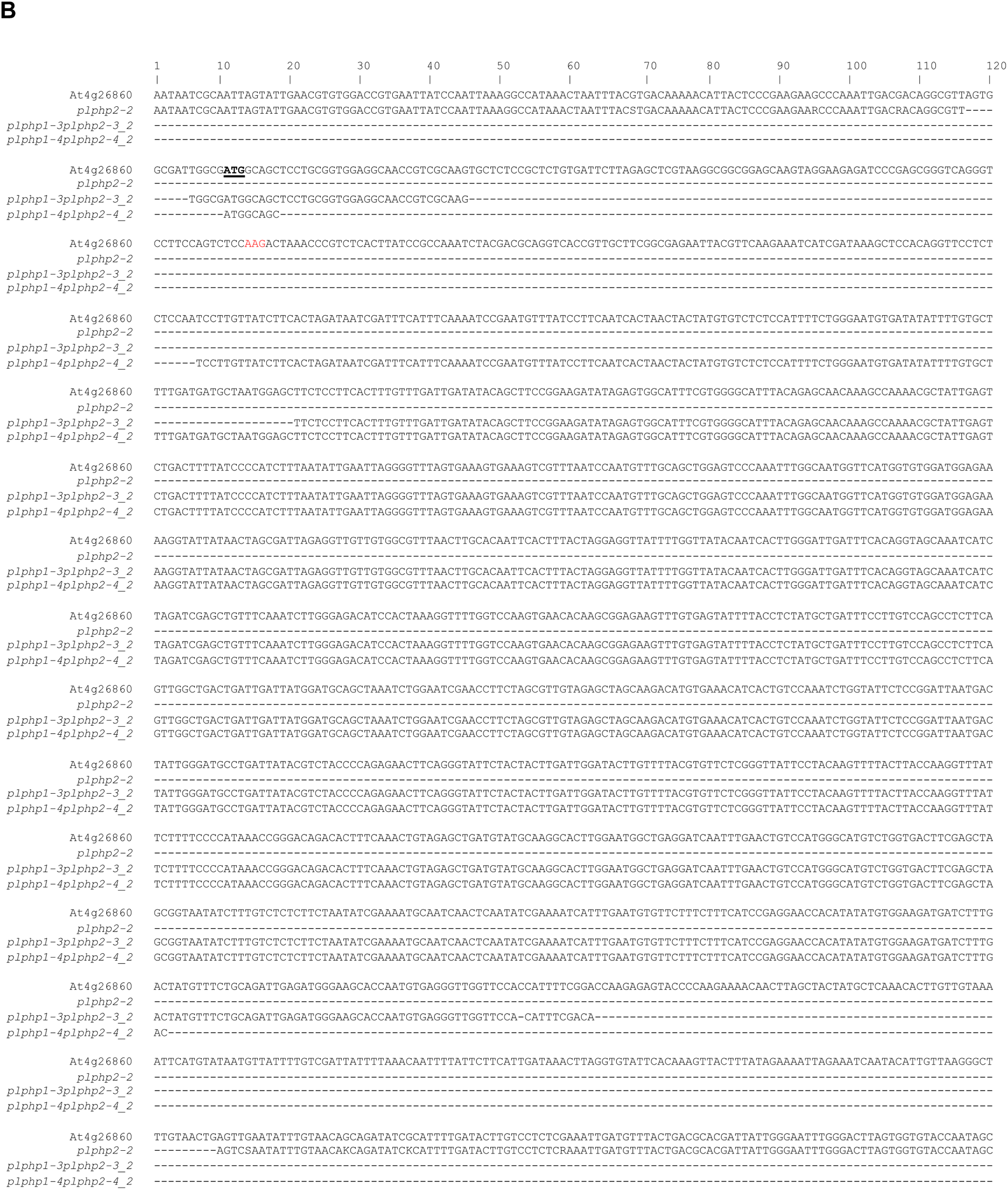
Sequencing analysis of loci At1g11930 (**A**) and At4g26860 (**B**) in lines generated by CRISPR/Cas9. The top row of the alignment shows the reference sequence of the respective loci. The translational start codon is underlined in bold, while the codon of the PLP binding lysine residue is highlighted in red at both loci. Genetic inversion in *plphp1-2* is indicated in blue and missing nucleotides by a dash.

**Supplementary Figure S3.**
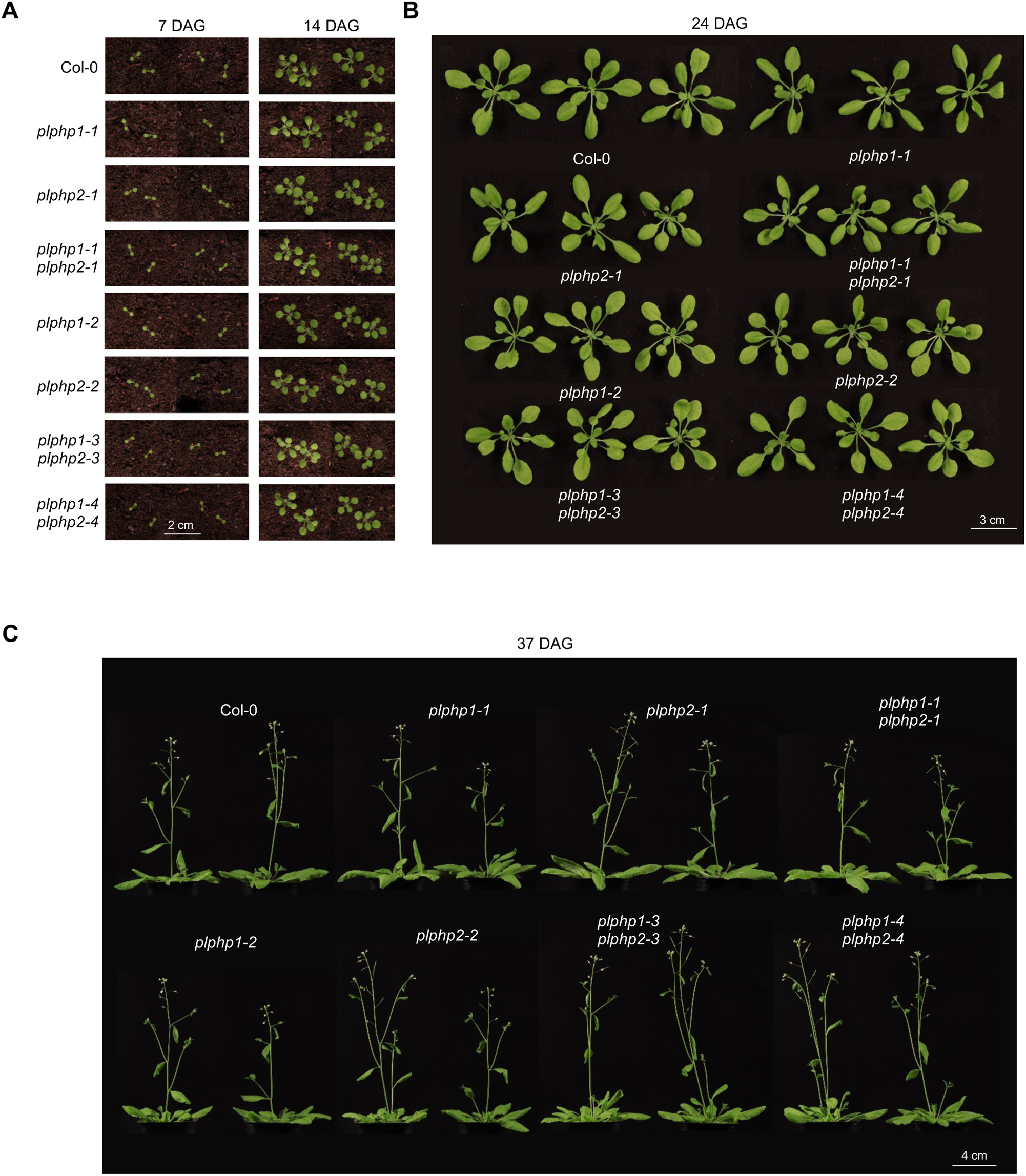
Morphological appearance of *plphp* lines compared to Col-0 (wild type). Aerial organ development of single and double mutants of *AtPLPHP1/2* compared to Col-0 throughout the life cycle of the plant. Photos were captured at the indicated days after germination (DAG). Plants were grown on soil under a 16-hour photoperiod at 120 μmol photons m^−2^ s^−1^ generated by fluorescent lamps at 22°C and 8-hour darkness at 18°C with 60-70% relative humidity and ambient CO_2_.

**Supplementary Figure S4.**
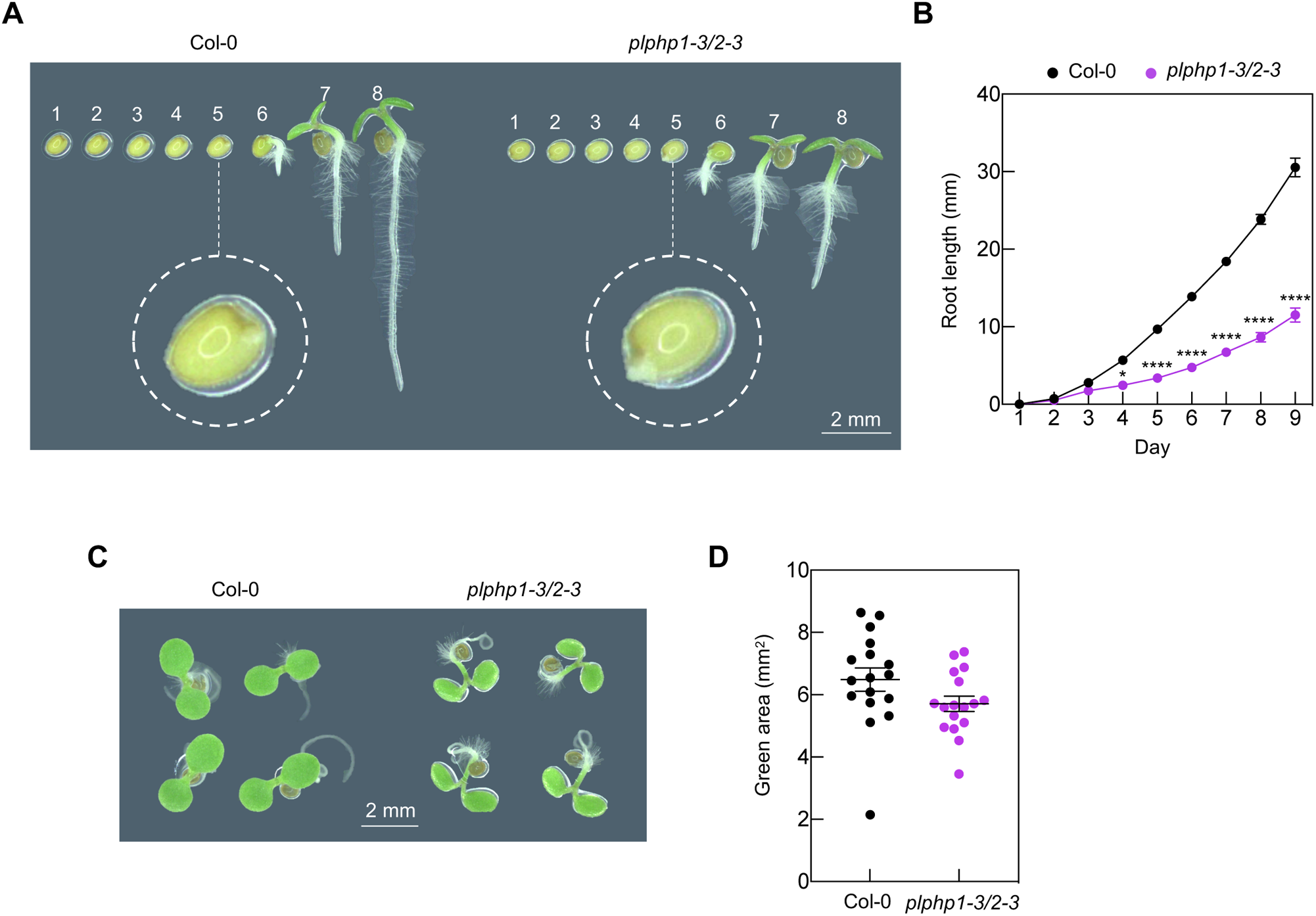
Detailed analysis of germination and early seedling development. **A)** Representative images of germination of Col-0 (wild type) and *plphp1-3/2-3*. The indicated numbers correspond to the following stages: 1: 1 hour after sowing, 2: 7 hours after sowing, 3: 0 hours after stratification, 4: 8 hours after stratification, 5: 26 hours after stratification, 6: 46 hours after stratification, 7: 70 hours after stratification, 8: 93 hours after stratification. The inset zooms in on the stage of radical emergence. **B)** Quantification of the root length of lines in **A** during germination, seedling establishment and early seedling development on vertical culture plates grown under a 16-hour photoperiod (120 μmol photons m^−2^ s^−1^) at 22°C and 8 hours dark at 18°C. Statistical analysis was performed by multiple *t-*tests (p*≤0.05, ****p≤0.0001). **C)** The hypocotyls of *plphp1-3/2-3* lean over and lay parallel to the surface of the media when grown on horizontal culture plates. Image taken 5 days after germination. **D)** Cotyledon area quantification of seedlings from **C**. Error bars indicate the standard error of the mean. An unpaired two-tailed *t-*test with Welch’s correction did not reveal significant differences between the means (p=0.0922).

**Supplementary Figure S5.**
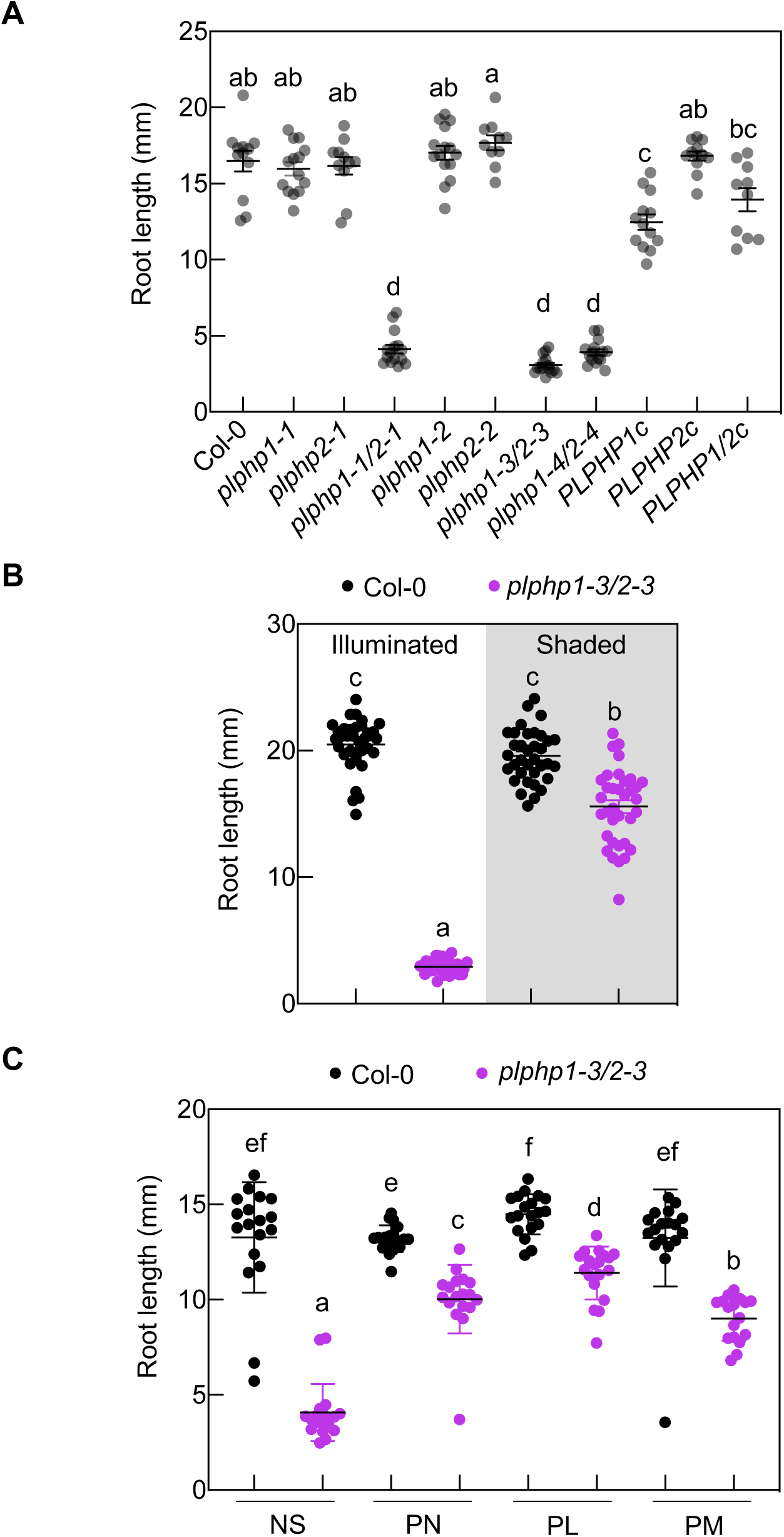
A) Root length of plant lines as indicated (Col-0 is the wild type; T-DNA insertion mutants *plphp1-1, plphp2-1*, and the cross *plphp1-1 plphp2-1*, CRISPR mutants *plphp1-2, plphp2-2*, *plphp1-3 plphp2-3*, *plphp1-4 plphp2-4*, as well as *plphp1-3/2-3* carrying either *PLPHP1* (*PLPHP1c*), *PLPHP2* (*PLPHP2c*), or both *PLPHP1* and *PLPHP2* (*PLPHP1/2c*), transgenes, respectively) grown with full illumination on vertical culture plates under a 16-hour photoperiod (120 μmol photons m^−2^ s^−1^) at 22°C and 8 hours dark at 18°C at 5 days after germination from Figure 3D. Error bars represent standard error of mean. Letters represent significantly different groups (p≤0.05) according to a Brown-Forsythe ANOVA test with Dunnett’s T3 multiple comparison. **B)** Root length of *plphp1-3/2-3* grown as in **A** but with roots either illuminated (white background) or shaded (gray background) compared to Col-0 from Figure 3E. **C)** Root length of *plphp1-3/2-3* grown as in **A** with roots illuminated and either without supplementation (NS) or supplemented with either pyridoxine (PN), pyridoxal (PL) or pyridoxamine (PM) (5 μM in each case) compared to Col-0, from Figure 5A.

**Figure S6.**
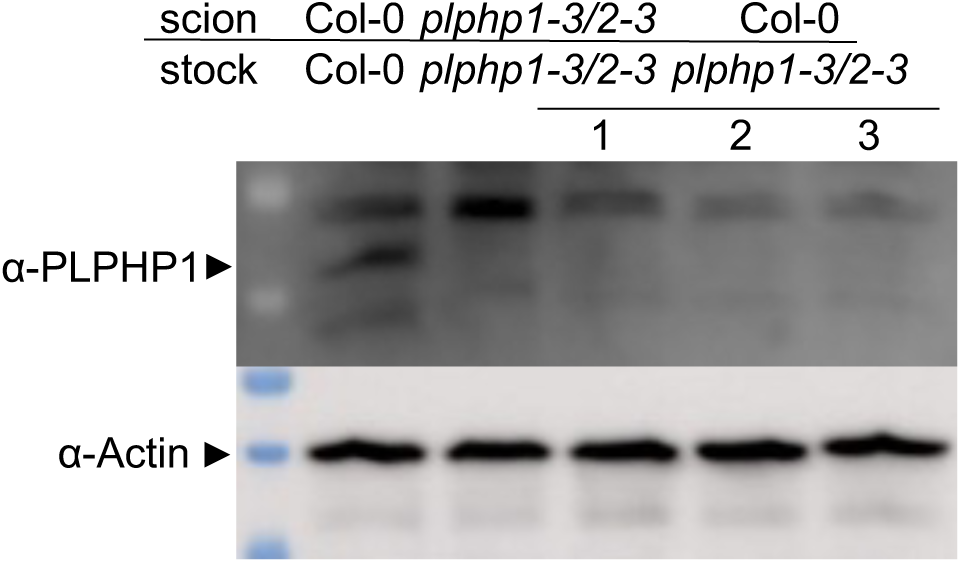
Immunochemical analysis of AtPLPHP1 in crude root extracts of micrografted seedlings of self-grafted Col-0, *plphp1-3/2-3* or grafts of Col-0 scions onto *plphp1-3/2-3* stocks as indicated. Seedlings were first grown with the root area shaded for six days to provide comparable root morphology at the time of grafting. Micrografting was then performed, and a recovery period of six days was allowed during which the root area was illuminated. After recovery, the grafts were transferred to rhizoboxes (Gorelova et al. 2022) and grown for a further 15 days during which the root area was illuminated. Protein (7.5 μg) was extracted from the roots and subjected to immunochemical analysis. The α-actin antibody on the same blot serves as a loading control.

**Figure S7.**
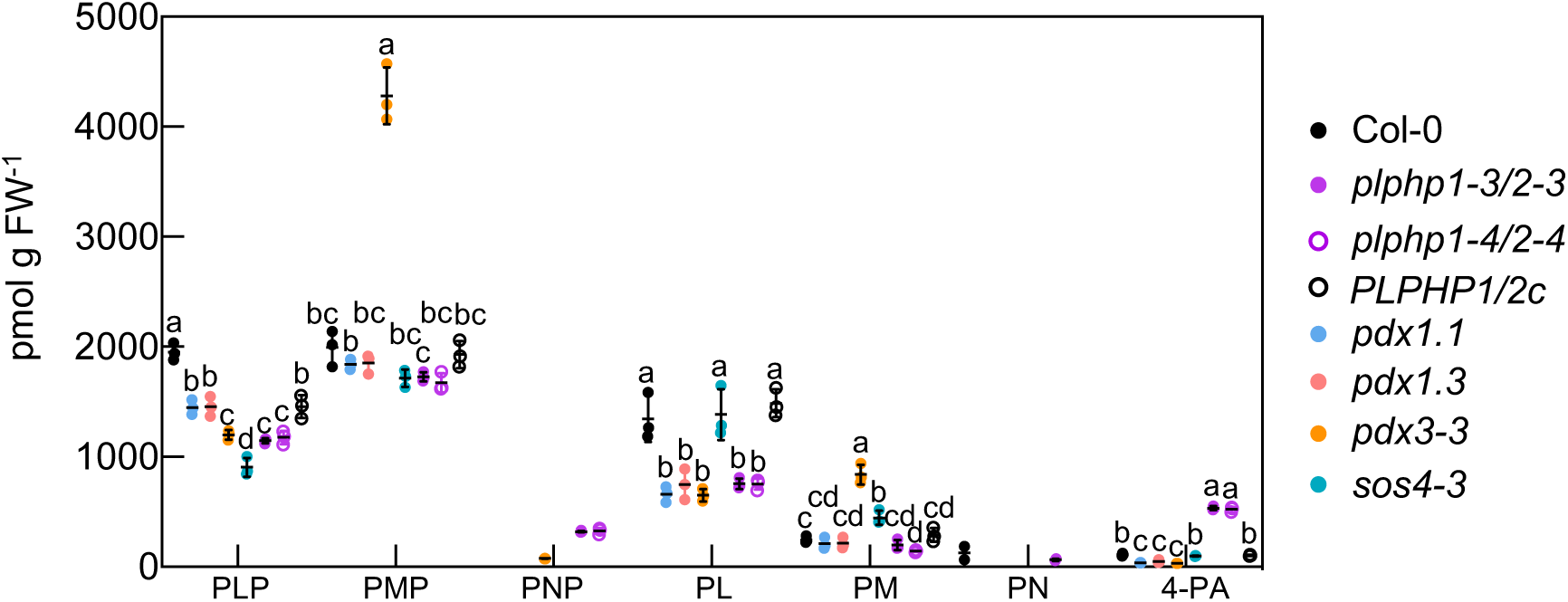
Scatter dot plot showing the abundance of vitamin B_6_ related compounds (pyridoxal 5ʹ-phosphate (PLP), pyridoxamine 5ʹ-phosphate (PMP), pyridoxine 5ʹ-phosphate (PNP), pyridoxal (PL), pyridoxamine (PM), pyridoxine (PN), 4-pyridoxic acid (4-PA)) in whole seedlings of the lines as indicated (Col-0 is the wild type; CRISPR mutants *plphp1-3 plphp2-3*, *plphp1-4 plphp2-4*; *plphp1-3/2-3* carrying both *PLPHP1* and *PLPHP2* (*PLPHP1/2c*) transgenes; *pdx1.1* and *pdx1.3* (biosynthesis *de novo* mutants); as well as *pdx3-3* and *sos4-3* (salvage pathway mutants) at 10 days after germination grown with full illumination on vertical culture plates under a 16-hour photoperiod (120 μmol photons m^−2^ s^−1^) at 22°C and 8 hours dark at 18°C. Letters represent significantly different groups discovered in a Brown-Forsythe ANOVA test with multiple *t*-tests with Welch’s correction (p≤0.05).

**Figure S8.**
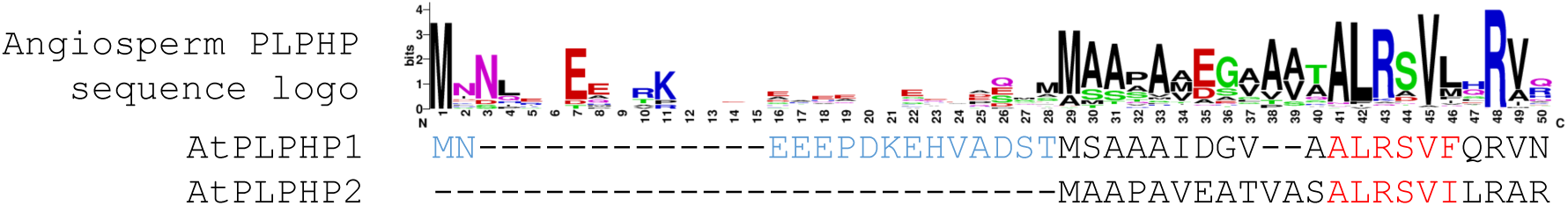
Predicted mitochondrial targeting sequence in PLPHPs. The first 50 residues of the Angiosperm PLPHP consensus is shown at the top as a sequence logo representing amino acid conservation at a given position and the corresponding sequences of AtPLPHP1 and AtPLPHP2 below. The key part of the mitochondrial targeting sequence predicted by MitoFates (Fukasawa et al., 2015) is highlighted in red with 0.581 targeting probability in AtPLPHP2 and 0.001 or 0.578 probability in AtPLPHP1 when the N-terminal extension (in blue) was included or omitted, respectively.

**Supplementary Table S1.** Distribution of PLPHP orthologs annotated in the Kyoto Encyclopedia of Genes and Genomes under the gene orthology identifier K06997 in various domains of life.

|  | Animals |  | Plants |  | Fungi |  | Protists |  | Bacteria |  | Archaea |  |  |  |
| --- | --- | --- | --- | --- | --- | --- | --- | --- | --- | --- | --- | --- | --- | --- |
|  | Occurrence |  | Occurrence |  | Occurrence |  | Occurrence |  | Occurrence |  | Occurrence |  |  |  |
| # ortholog | # taxa | % | # taxa | % | # taxa | % | # taxa | % | # taxa | % | # taxa | % |  |  |
| 1 | 504 | 92.31 | 26 | 18.31 | 142 | 98.61 | 49 | 94.23 | 6555 | 95.33 | 33 | 100 |  |  |
| 2 | 40 | 7.33 | 81 | 57.04 | 2 | 1.39 | 3 | 5.77 | 314 | 4.57 | 0 | 0 |  |  |
| 3 | 2 | 0.37 | 19 | 13.38 | 0 | 0 | 0 | 0 | 7 | 0.10 | 0 | 0 |  |  |
| 4 | 0 | 0 | 11 | 7.75 | 0 | 0 | 0 | 0 | 0 | 0 | 0 | 0 |  |  |
| 5 | 0 | 0 | 2 | 1.41 | 0 | 0 | 0 | 0 | 0 | 0 | 0 | 0 |  |  |
| 6 | 0 | 0 | 3 | 2.11 | 0 | 0 | 0 | 0 | 0 | 0 | 0 | 0 |  |  |
| 7 | 0 | 0 | 0 | 0 | 0 | 0 | 0 | 0 | 0 | 0 | 0 | 0 |  |  |
|  | 546 taxa |  | 142 taxa |  | 144 taxa |  | 52 taxa |  | 6876 taxa |  | 33 taxa |  | SUM taxa | 7793 |
|  | 590 orthologs |  | 317 orthologs |  | 146 orthologs |  | 55 orthologs |  | 7204 orthologs |  | 33 orthologs |  | SUM orthologs | 8345 |

**Supplementary Table S2.**
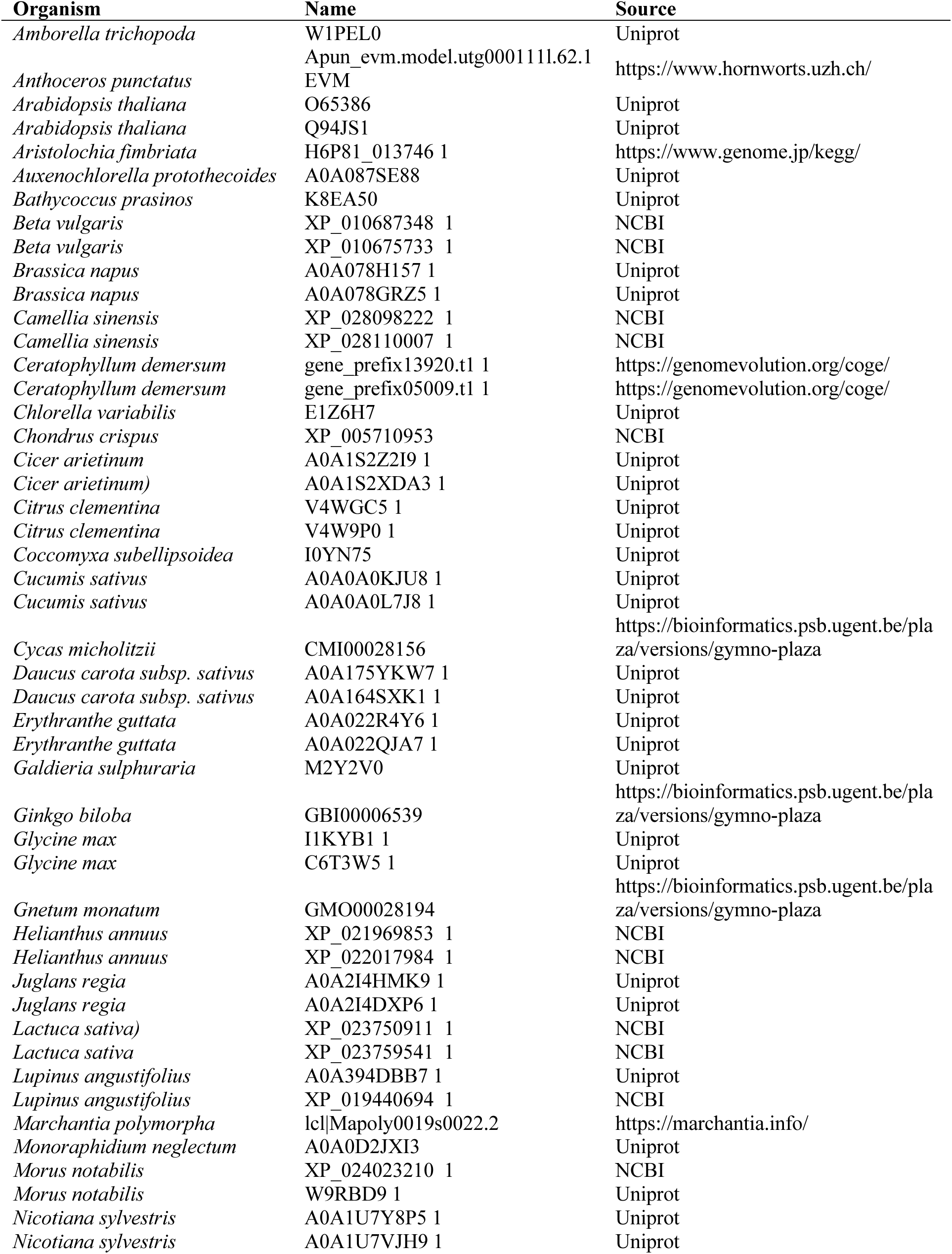
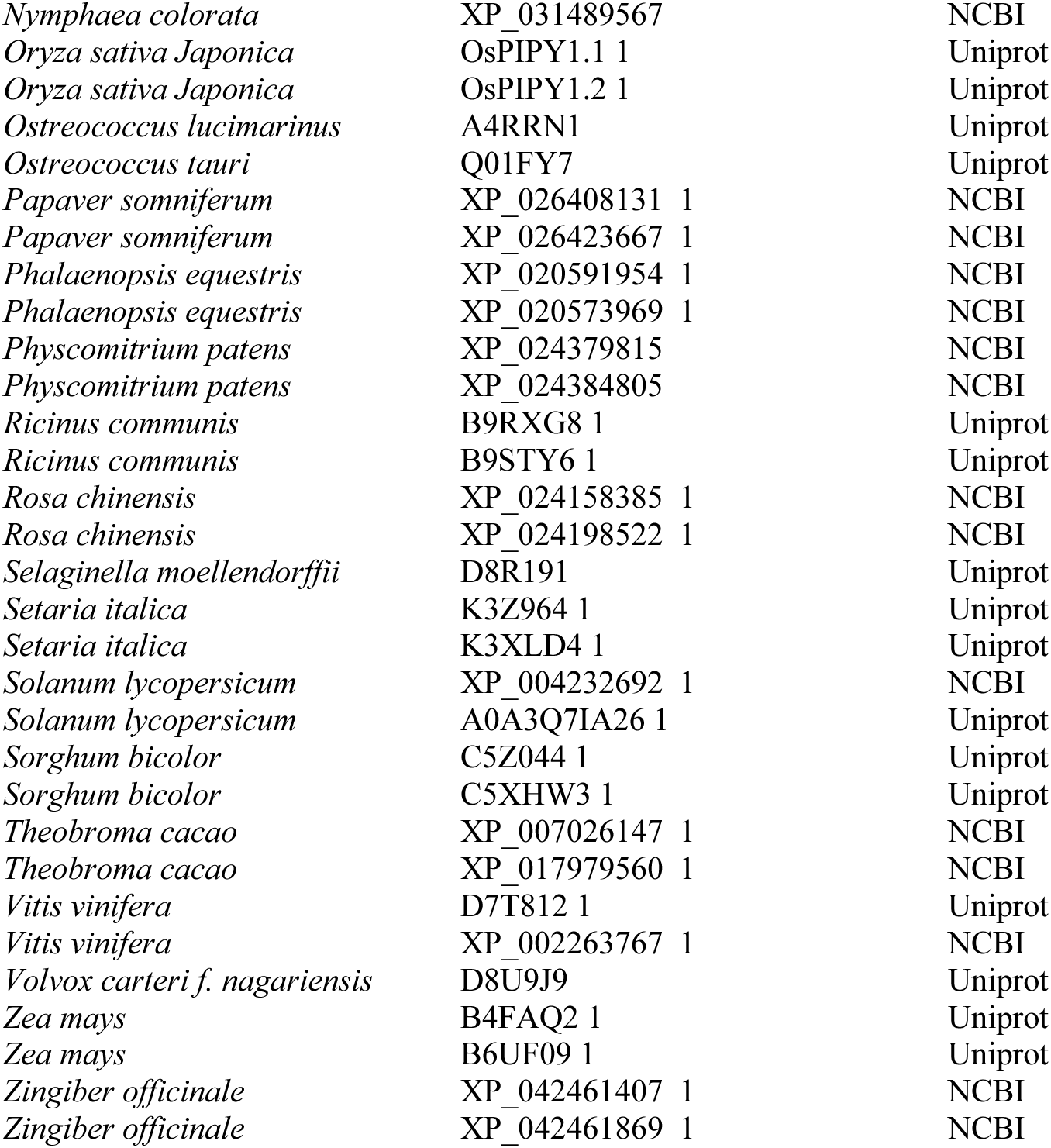
Representative PLPHP orthologs within Archaeplastida from Figure 1A.

**Supplementary Table S3.** List of lines generated and used in this study.

| Name | Mutagen | Gene | Locus | Nature of mutation | Region affected | Background | Source |
| --- | --- | --- | --- | --- | --- | --- | --- |
| *plphp1-1* | T-DNA | *PLPHP1* | At1g11930 | SAIL_675_A03 | 1351 (exon 6) | Col-0 | NASC |
| *plphp2-1* | T-DNA | *PLPHP2* | At4g26860 | GK_740_A10 | 21 (exon 1) | Col-0 | NASC |
| *plphp1-1/2-1* | T-DNA | *PLPHP1* | At1g11930 | SAIL_675_A03 | 1351 (exon 6) | Col-0 | this |
|  |  | *PLPHP2* | At4g26860 | GK_740_A10 | 21 (exon 1) |  | study |
| *plphp1-2* | CRISPR | *PLPHP1* | At1g11930 | reversion (106-261), Δ261-1057 | Exon 1-5 | Col-0 | this study |
| *plphp2-2* | CRISPR | *PLPHP2* | At4g26860 | $\Delta$(-)12-1561 | Whole CDS | Col-0 | this study |
| *plphp1-3/2-3* | CRISPR | *PLPHP1* | At1g11930 | Δ106-261 | Exon 1 | Col-0 | this |
| | | *PLPHP2* | At4g26860 | Δ35-370 | Exon 1 | | study |
| *plphp1-4/2-4* | CRISPR | *PLPHP1* | At1g11930 | Δ261-573, +8 pb | Exon 2-3 | Col-0 | this |
| | | *PLPHP2* | At4g26860 | Δ8-236 | Exon 1 | | study |

| Plant line | Type | Background | Clone | Gene | Promoter | Source |
| --- | --- | --- | --- | --- | --- | --- |
| *PLPHP1c* | complementing | *plphp1-3/2-3* | (-)1533-1830 of PLPHP1 | *PLPHP1* | endogenous | this study |
| *PLPHP2c* | complementing | *plphp1-3/2-3* | (-)683-1660 of PLPHP2 | *PLPHP2* | endogenous | this study |
| *PLPHP1/2c1* | complementing | *plphp1-3/2-3* | (-)1533-1830 of PLPHP1 | *PLPHP2* | endogenous | this study |
|  |  |  | (-)683-1660 of PLPHP2 |  |  |  |
| *PLPHP1_K55A* | Lys mutant | *plphp1-3/2-3* | PLPHP1_K55A<br>in *PLPHP1c* template | *PLPHP1* | endogenous | this study |
| *PLPHP2_K42A* | Lys mutant | *plphp1-3/2-3* | PLPHP2_K42A<br>in *PLPHP2c* template | *PLPHP2* | endogenous | this study |

**Supplementary Table S4.**
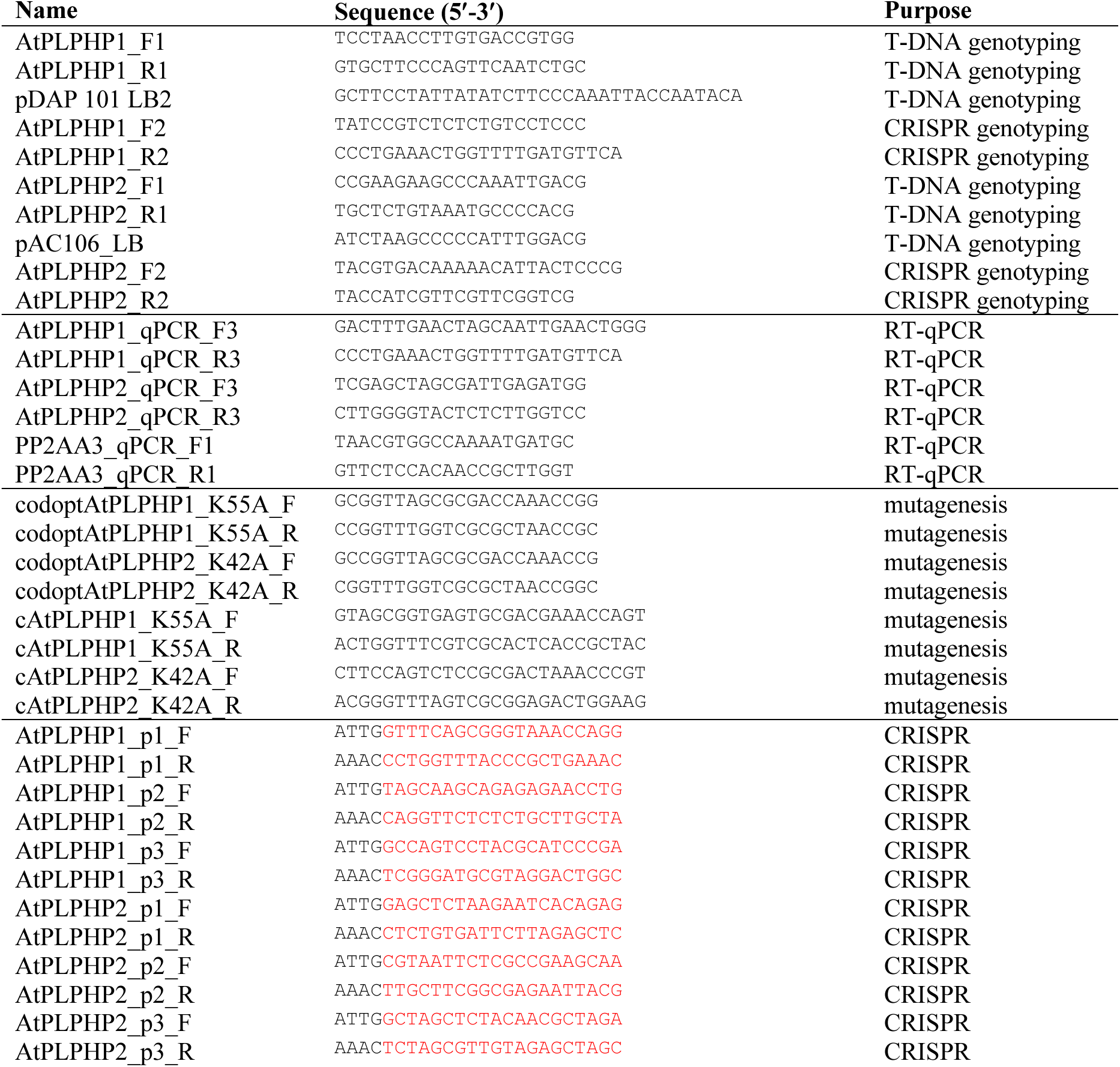
List of oligonucleotides used in this study. pDAP_101_LB2 and pAC106_LB are specific for the left border sequence of the *pDAP_101* and *pAC106_LB* plasmid T-DNA regions in *plphp1-1* and *plphp2-1*, respectively. Oligonucleotides used to generate mutations in constructs codon optimized for recombinant protein expression in *Escherichia coli* are marked as “codopt”. Protospacers are indicated in red.

